# Multi-species atlas resolves an axolotl limb development and regeneration paradox

**DOI:** 10.1101/2023.03.01.530572

**Authors:** Jixing Zhong, Rita Aires, Georgios Tsissios, Evangelia Skoufa, Kerstin Brandt, Tatiana Sandoval-Guzmán, Can Aztekin

## Abstract

Humans and other tetrapods are considered to require apical-ectodermal-ridge, AER, cells for limb development, and AER-like cells are suggested to be re-formed to initiate limb regeneration. Paradoxically, the presence of AER in the axolotl, the primary regeneration model organism, remains controversial. Here, by leveraging a single-cell transcriptomics-based multi-species atlas, composed of axolotl, human, mouse, chicken, and frog cells, we first established that axolotls contain cells with AER characteristics. Surprisingly, further analyses and spatial transcriptomics revealed that axolotl limbs do not fully re-form AER cells during regeneration. Moreover, the axolotl mesoderm displays part of the AER machinery, revealing a novel program for limb (re)growth. These results clarify the debate about the axolotl AER and the extent to which the limb developmental program is recapitulated during regeneration.

## Introduction

Vertebrate limb development requires apical-ectodermal-ridge, AER, cells at the dorsal-ventral boundary of developing limbs, which enable the expansion of limb bud mesodermal cells and provide patterning cues^1^. The AER supplies critical signaling ligands and its spatial organization contribute to the morphogen gradients to form a correctly patterned limb^2^. Interestingly, the leading limb regeneration model organism, the axolotl^3^, was suggested not to have an AER, as they lack a morphological ridge structure and some of the molecular AER markers^4–7^. Despite these findings, as regeneration is thought to largely recapitulate development, salamander regeneration is considered to re-form AER-like cells to act as a signaling center apical-epithelial-cap, AEC, at the amputation plane^8–10^. The AEC is required to form a connective-tissue lineage-rich blastema for regeneration^11, 12^, and its absence in mammals has been long-hypothesized to be one of the reasons for the regeneration-incompetency^13, 14^. However, previous reports provided conflicting results for the AER and AEC marker expressions^4, 7, 15–21^, and a sub group of the axolotl connective tissue (CT) cells was suggested to express some of the AER-related genes^4, 7, 21, 22^. Consequently, different conclusions have been drawn based either on morphological assessments or assaying the expression of a small set of specific marker genes. Because of this, the existence of AER in axolotls and the re-use of AER-like cells for salamander limb regeneration remain unclear. Unbiased and comprehensive analyses are warranted to resolve the cellular identity of this critical population in the contexts of limb development, regeneration, and evolution studies.

### Multi-species limb atlas is generated by integrating limb datasets from five vertebrates

To determine if the developing axolotl limbs have an AER-like population, we aimed to benefit from a single-cell transcriptomics-based multi-species limb atlas (**Figure 1A)**. We hypothesized that if AER-like cells are present in the axolotl limb, their molecular identity would overlap with the AER cells from other species. To test this, we first collected publicly available scRNA-Seq datasets of developing limbs from axolotls (*Ambystoma mexicanum*)^23^ and, species with their developmental stages that are documented to have an AER: humans (*Homo sapiens*)^24, 25^, mice (*Mus musculus*)^26, 27^, chickens (*Gallus gallus*)^28^, and frogs (*Xenopus laevis*)^23, 29^) (**Figures S1 and S2 and Supplementary Table 1)**. Then, using Seurat integration, which has been shown to integrate cross-species datasets with high accuracy^30^, we established a multi-species limb atlas that in total contains 50,248 representative cells from all samples (**Figures 1B and S3**). Coarse lineage annotation detected various mesodermal and ectodermal populations in the multi-species limb atlas, and finer annotation captured the AER cluster (**Figures 1C and S3**). Overall, the multi-species atlas was able to group cells based on their potential cell identity rather than the species or the experiment batch (**Figure S3B**).

**Figure 1.**
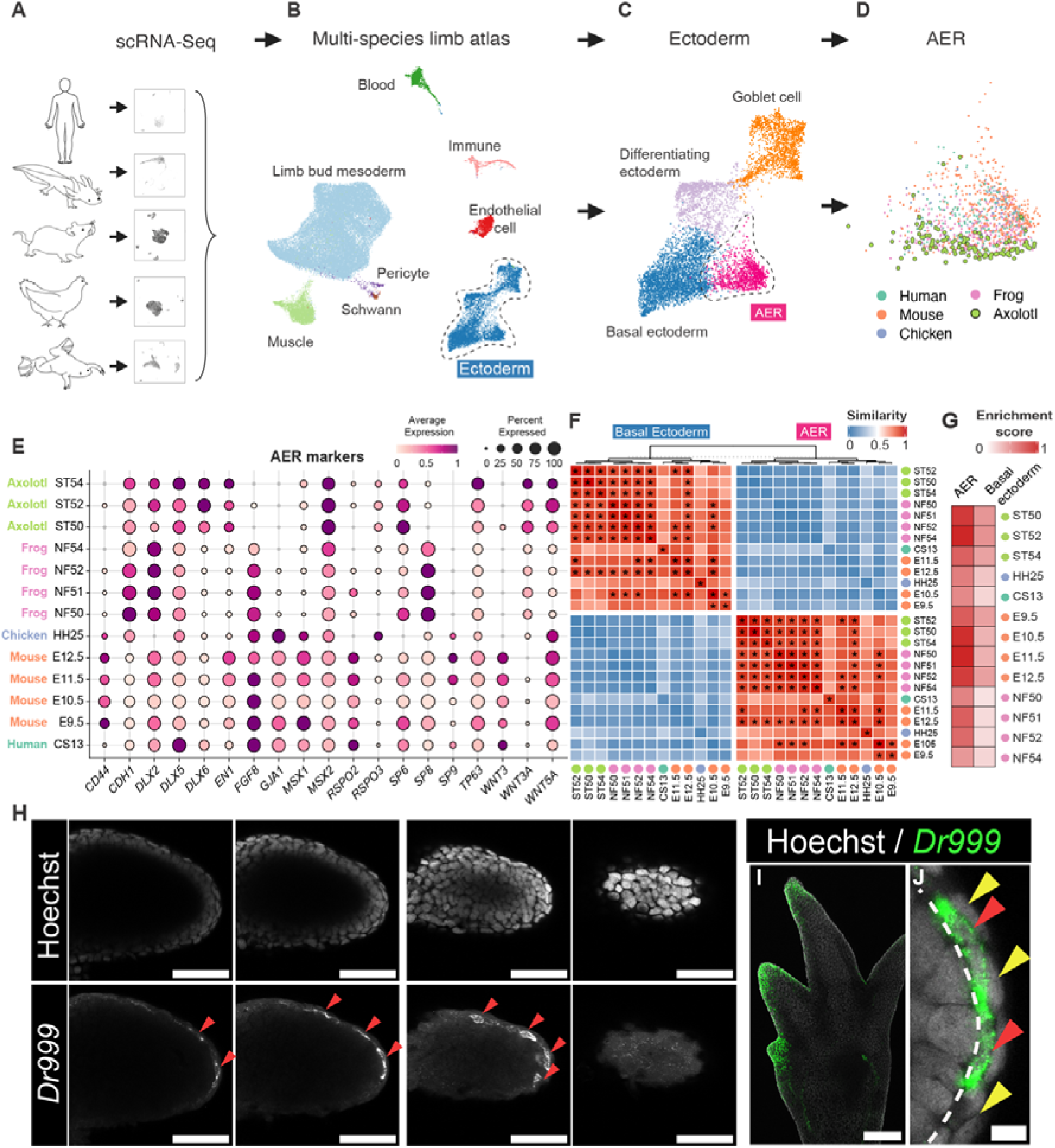
Multi-species limb atlas reveals developing axolotl limbs have cells with apical-ectodermal-ridge-(AER) characteristics. A. Analyzed species and examples of the used publicly available single-cell RNA-Seq (scRNA-Seq) datasets (please see Figure S1 and Table 1 for the full list) to generate a multi-species limb atlas are illustrated. B. UMAP plot of Seurat-integrated multi-species limb atlas is shown. Individual datasets from each species with different developmental stages (please see Figure S1 and Table 1 for the full list) are integrated. Clustering and annotation based on marker gene expressions (please see Figure S3) are used to label lineage and cell type identities, indicated by different colors and text. C. UMAP plot of subclustering of the basal ectodermal lineage of multi-species limb atlas is shown. Subclustered cell identities are based on marker genes that are listed in Figure S3, and labeled by different colors and text. D. UMAP plot of species contribution to the apical-ectodermal-ridge (AER) cluster of multi-species limb atlas is visualized. Cells from different species are color-coded. E. Dot plot showing AER marker expressions in AER cells from different species. Names of the species are color-coded, and developmental stages are indicated. The dot color indicates the mean expression that was normalized to the max of each dataset and to the max of each gene; the dot size represents the percentage of cells with non-zero expression. F. Heatmap showing the MetaNeighbor score for pair-wise similarities of basal ectoderm and AER clusters across species and different developmental stages. X and Y axes indicate different species and developmental stages. Species are color-coded, and developmental stages are indicated in the text. Asterisks (*) denote the pairs with similarity scores above 0.9. G. Heatmap showing signaling ligands (see Table S2 for the gene set) gene set enrichment analysis scores for AER, and basal ectoderm clusters for each species. The basal ectoderm represents the transcriptome-wide most similar population to the AER, and is used in comparison to highlight specific enrichment in the AER cluster. Colored dots in the Y axis indicate different species, and developmental stages are indicated in the text. H. Single optical section of z-stacks of confocal images of Stage 46 axolotl forelimb buds stained for *Dr999-Pmt21178* mRNA via hybridization-chain-reaction are shown. Different z-stacks were shown from left to right, representing different levels of the dorsal-ventral axis. (Top) Gray, Hoechst; Bottom Gray, *Dr999-Pmt21178* mRNA *(hereafter, referred to as Dr999 in the figure)*. Scale bar: 100 μm. I. Max-projection confocal image of Stage 53 axolotl forelimb digit tips stained for *Dr999-Pmt21178* mRNA via HCR. Green, *Dr999-Pmt21178* mRNA; Gray, Hoechst. Scale bar: 250 μm. J. Zoomed single optical section image of the axolotl limb bud from (H) stained for *Dr999-Pmt21178* mRNA. Red arrows show *Dr999-Pmt21178* + squamous cells, and yellow arrows show outer layer peridermal cells. The basement membrane is labeled with a dashed line. Green, *Dr999-PMT21178* mRNA; Gray, Hoechst. Scale bar: 10 μm.

### Developing axolotl limbs contain cells with apical-ectodermal-ridge (AER) features

Examining the species contribution to the multi-species AER cluster, we found cells from developing axolotl limbs **(Figure 1D)**. A second cross-species data integration approach, SAMap^31^, yielded the same result, suggesting that axolotl limb buds contain AER-like cells regardless of the integration method (**Figure S4**). Then, we surveyed the established AER markers to determine the transcriptional similarities of these axolotl cells with the AER in other species (**Figures 1E and S5**). The large majority of axolotl cells in this cluster co-expressed many of the AER markers^1^, such as *Wnt5a* and *Msx2*, whilst some others (e.g., *Fgf8*) were absent **(Figures 1E and S5)**, in alignment with recent reports^4, 7^. A further transcriptome-wide comparison found not only a high similarity of AER-like axolotl cells to the AER cells in other species, but also that this population was distinct from non-AER basal ectodermal cells **(Figures 1F and S6)**. Hence, our results provide high-throughput evidence that developing axolotl limbs contain cells with AER transcriptional programs as in other species, and these cells are hereafter referred to as axolotl AER cells.

Next, we evaluated the potential functional properties of the axolotl AER. As the AER is a well-recognized signaling center^1^, we aimed to reveal if they express ligands from critical limb development-related signaling pathways. We generated a potential secretome gene group, composed of ligands from FGFs, BMPs, WNTs, NOTCH/DELTAs, and TGFbs **(Supplementary Table 2)**, and performed gene set enrichment analysis on cell clusters. Axolotl AER cells had comparable signaling potential to the AER in other species **(Figures 1G and S7)**. Meanwhile, individual pathway examination revealed the axolotl AER differs in paralog and co-factor expressions, particularly in the FGF and WNT pathways **(Figure S8)**, complementing and extending previous observations^4, 7, 21^. We could not detect noticeable differences in transcriptional levels associated with DELTAs, BMPs, or TGFbs **(Figure S8)**. Overall, axolotl AER cells have comparable transcriptional levels for limb development-related signaling pathway ligands, albeit potentially critical differences remain.

The AER in other species forms at the dorsal-ventral boundary of developing limbs, which is considered to be critical in setting morphogen gradients for subsequent growth and patterning^2^. To determine if the axolotl AER cells have a similar spatial organization, we sought to visualize them. Using the scRNA-seq dataset, we identified *Dr999-Pmt21178* and *Vwa2* as the marker genes with high and specific expression in the axolotl AER cells **(Figures S9A and S9B)**. Whole-mount hybridization chain reaction on developing axolotl limbs from different stages demonstrated that these potential marker genes show specific expression at the dorsal-ventral boundary at the limb bud stages, albeit more scattered compared to AER localization in other species, **(Figures 1H and S9C)**, and digit tips during digit forming stages **(Figures 1I and S9C)**, resembling the spatial organization of AER in other species^1, 2^.

Next, we investigated the AER cellular morphology. Unlike amniotic or frog AER where AER cells have mainly cuboidal or columnar cell shape^1, 29^, we found that *Dr999-Pmt21178* positive axolotl AER cells mostly present a squamous shape **(Figures 1J and S9B)**. Moreover, these AER cells have a high degree of similarity to the outer skin cells, the periderm **(Figures 1J and S9B)**, which may explain why prior morphology-based studies could not distinguish them from other populations. Altogether, our imaging results highlight that the axolotl AER cells have a spatial organization in developing limbs similar to other species with a unique cellular morphology.

### Axolotl AER cells are not entirely re-formed during limb regeneration

Earlier studies suggested that the AER is re-formed during limb regeneration across different species acting as the AEC, based on morphological examination and a set of gene expression similarities with other species^8, 9^. In alignment with this proposition, previously, we found that at the single-cell level, limb regeneration-competent *Xenopus laevis* tadpoles re-use AER cell transcriptional program to act as the AEC^29^ **(Figure S10)**. Having identified axolotl limb buds containing cells with an AER transcriptional program, we asked if this program is re-used in the course of axolotl regeneration, as in frog tadpole regeneration.

To test whether the axolotl AER program is re-used during regeneration, we first combined publicly-available comprehensive and time-course axolotl limb regeneration (**Figure S11)** ^19^ and development **(Figure S2E)**^23^ datasets **(Figure 2A)**. We then focused on all the basal ectodermal cells, which would contain the AER or AEC cells. Subclustering of these basal ectodermal cells revealed that the axolotl AER cells gathered with some cells from the regeneration samples **(Figures 2B and 2C).** Investigation of the expression profile of these cells from the regeneration samples revealed highly specific expression of the known axolotl AEC markers (e.g., *Mdk*^12^*, Frem2*^18^*, Krt5*^32^) **(Figure S11D),** indicating significant similarity between the AER and the AEC. Nevertheless, we found that these AER and AEC cells have different localizations within the same cluster **(Figure 2C),** highlighting the potential transcriptional differences between them. Indeed, the comparison of AER and AEC cells by differentially expressed gene analysis identified significant transcriptional changes **(Figures 2D and 2E and S12 and Supplementary Table 3)**, some of which are specific to post-amputation samples (e.g., *Mmp13*) **(Figures 2D and 2E and S12)**.

**Figure 2.**
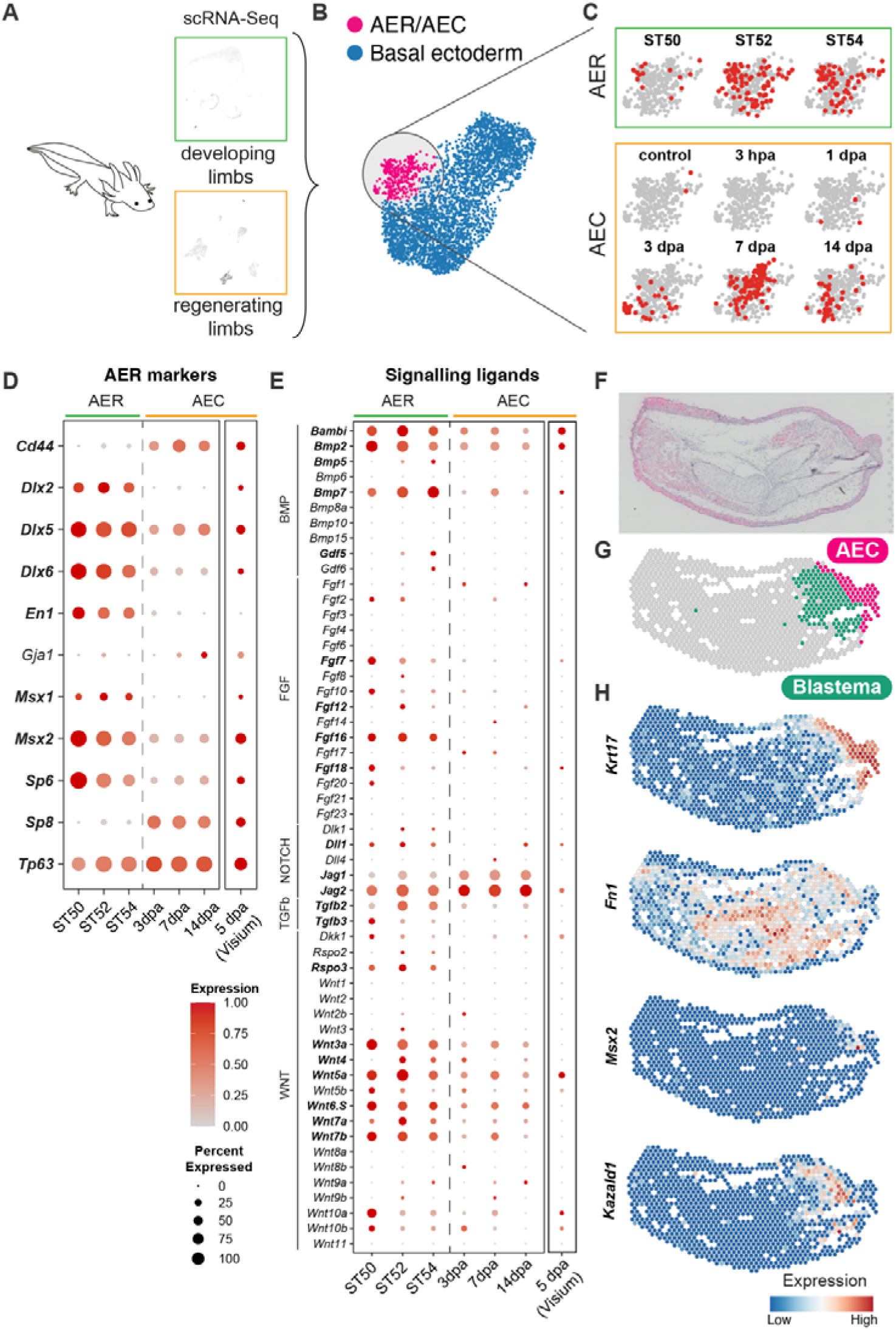
Axolotl AER cells are not entirely re-formed during limb regeneration. A. Schematics describing examples of the used scRNA-Seq datasets of axolotl limb development and regeneration are illustrated. B. UMAP plot of the basal ectoderm cells of the integrated axolotl limb development and regeneration datasets. Subclustered cell identities are labeled by different colors and text. Pink dots, AER or apical-epithelial-cap (AEC) cells; Blue dots, basal ectoderm cells. C. Sample contribution to integrated AER/AEC cluster from (B) is visualized. Red dots indicate cells from the selected sample; gray dots indicate the other cells in the AER/AEC cluster. hpa: hours-post amputation; dpa: days-post amputation. D. Dot plot showing AER marker expressions in (left) the scRNA-seq datasets of axolotl limb development and regeneration AER or AEC clusters, respectively, and (right) spatial transcriptomics (Visium) AEC cluster (G). The dot color indicates the mean expression that was normalized to the max of each gene; the dot size represents the percentage of cells with non-zero expressions. Please note that the Visium and scRNA-seq datasets were normalized separately within the dataset. Significant differentially expressed genes between AER and AEC were labeled in bold (two-sided Wilcoxon rank-sum test; P-values < 0.05). E. Dot plot showing signaling ligand expressions in (left) scRNA-seq datasets of axolotl limb development and regeneration AER or AEC cells, respectively, and (right) spatial transcriptomics (Visium) AEC cluster (G). The dot color indicates the mean expression that was normalized to the max of each gene; the dot size represents the percentage of cells with non-zero expressions. Please note that the Visium and scRNA-seq datasets were normalized separately within the dataset. Significant differentially expressed genes between AER and AEC were labeled in bold (two-sided Wilcoxon rank-sum test; P-values < 0.05). F. 5 dpa axolotl limb regeneration tissue section that is used for 10X Visium spatial transcriptomics (refer to as Visium) is shown and stained for hematoxylin and eosin. The tissue is oriented with the anterior to the top, the posterior to the bottom, the proximal to the left and the distal to the right. G. Clustering of the Visium spots identified known tissue types, including the AEC and the blastema, labeled by pink and green colors, respectively. Please see Figure S13 for the full clustering results. H. Expression profiles of selected markers in the AEC and the blastema clusters are visualized.

Then, we asked if we could detect comparable signaling properties in the axolotl AEC to the axolotl AER cells. Surprisingly, we failed to identify high levels of expression for some of the ligands belonging to the mainly studied signaling pathways in this population, which contrasted with their developmental counterparts (**Figure 2E)**. Specifically, in axolotl AEC cells there was a lack of FGF pathway related ligands and quantitative expression differences for other ligands **(Figure 2E)**.

To validate these results and eliminate the possibility that scRNA-Seq did not capture cells specifically from the amputation plane, we performed spatial transcriptomics on a regenerating limb where the AEC and the blastema were evident **(Figures 2F-2H and S13)**. With this data we could identify the spatial distribution of the expected tissues types, including the AEC cluster located at the tip of the amputated limbs, as well as the blastema **(Figures 2G and 2H and S13)**, both of which were previously difficult to pinpoint using the conventional scRNA-Seq approach in axolotls. Nonetheless, when evaluating this AEC cluster, we could not detect certain AER marker genes and high ligand expressions again **(Figures 2D and 2E)**. Altogether, our results suggest the axolotl AEC is distinct from the axolotl AER developmental program. Critically, it lacks a high level of expression for developmental signaling ligands, unlike in frog tadpoles, emphasizing different regenerative programs between these two species.

### Mesodermal cells showing part of the epithelial AER transcriptional program are found in axolotl limbs

Interestingly, a few of the AER-associated genes, such as *Fgf8,* were reported to be expressed in the salamander anterior mesoderm^4, 7, 22^, raising the possibility that axolotl mesodermal lineage might show features of the signaling center epithelial AER transcriptional program. Beyond single-gene investigations, we sought to comprehensively evaluate this possibility.

To test this, we leveraged our multi-species limb atlas, which underscored a transcriptional program related to AER cell identity across analyzed species **(Figures 1C and 1D and S3 and S4)**. First, we identified the differentially expressed genes in the AER cluster in the atlas **(Supplementary Table 4)**. Second, we used consensus non-negative matrix factorization (cNMF) to detect transcriptional modules related to cell-identity and cell-activity within specific populations^33^ **(Figure S14 and Supplementary Table 4)**. Then, using these AER-related gene sets, we surveyed the expression pattern in mesodermal populations, including using gene set enrichment analysis **(Figure S15)**. In parallel, we also tested whether clustering based on these gene sets would aggregate AER-related ectodermal cells and mesodermal cells together, indicative of a high degree of similarity **(Figures 3A-3C and S16-S20)**.

**Figure 3.**
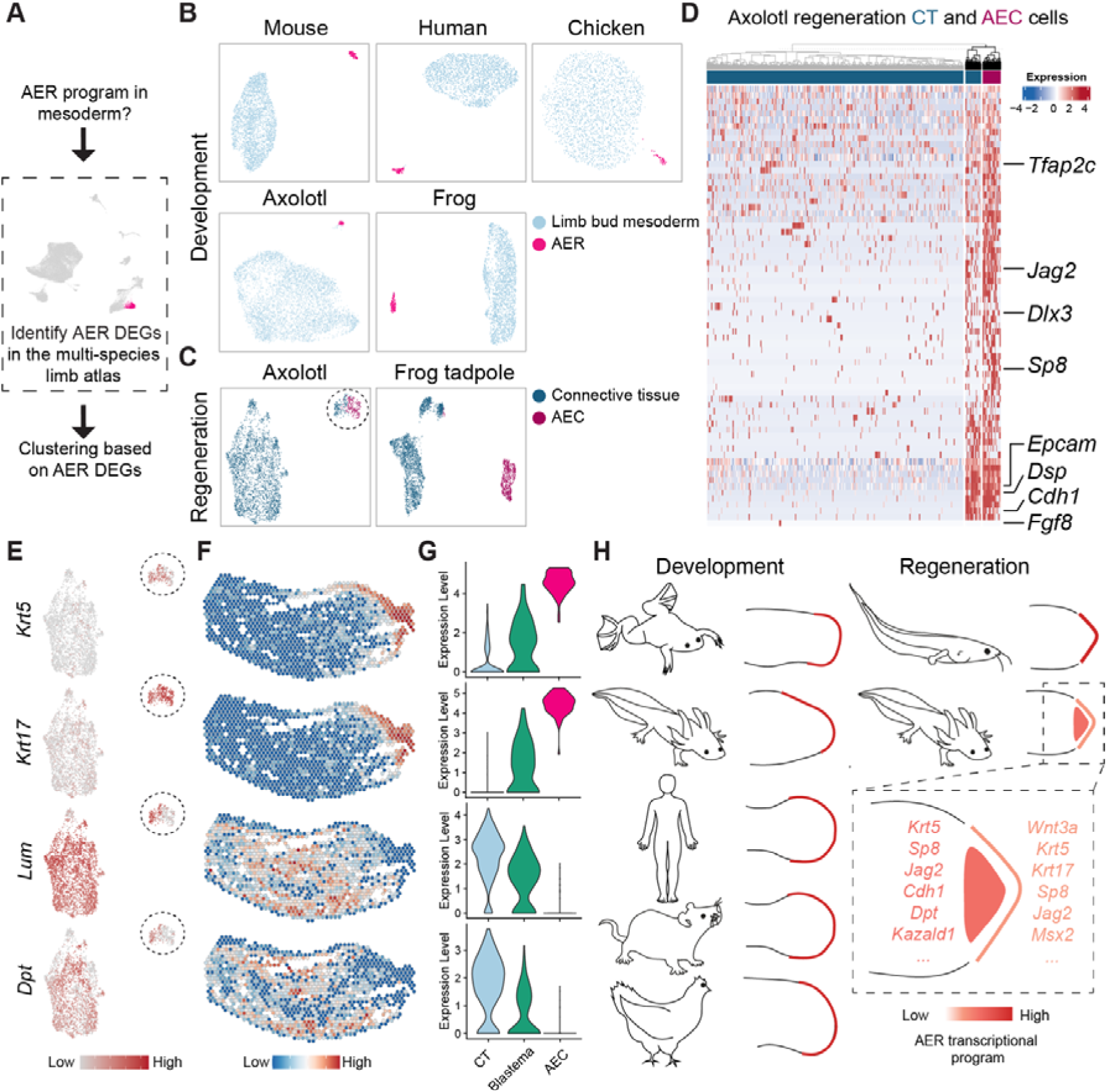
Axolotl limbs contain mesodermal cells showing part of the epithelial AER transcriptional program. A. Schematics of the strategy to evaluate the mesodermal cells showing the AER transcriptional program are shown. AEC and connective tissue (CT) cells are clustered based on differentially expressed genes in the AER cluster of the multi-species limb atlas. B. UMAP plots of the clustering based on the differentially expressed genes in the AER cluster of the multi-species limb atlas (Figure 1B) are shown for development datasets. Mouse E10.5, chicken E4.5, human CS13, frog ST51, and axolotl ST52 UMAPs are shown, and the full data with other developmental stages are presented in Figure S15. Analyzed animals are labeled. Light blue, limb bud mesoderm cells; pink, AER cells. C. UMAP plots of the clustering based on the differentially expressed genes in the AER cluster of the multi-species limb atlas are shown for the axolotl and frog tadpole regeneration datasets. Please note that these figures are the same as in Figure S15. Analyzed animals are labeled. Dark blue, CT cells; dark pink, AEC cells. CT cells gathered with the AEC and the AEC are circled with a dashed line. D. Heatmap showing shared gene expressions in the axolotl CT and AEC populations. Dendrogram based on gene expression profile indicates a subset of CT cells shows similarity to the AEC population, and is highlighted with black dendrograms. Note that these shared genes were identified by K-means clustering (Figure S20A) of the whole AER gene list used in Figures 3B and 3C., except for the manually added *Fgf8* due to its high relevance to AER. E. The expression profile of the example epithelial and fibroblast genes in the UMAP plot in Figure 3C axolotl regeneration dataset. F. Expression profiles of the example epithelial and fibroblast genes are visualized on the Visium dataset. Please note that the *Krt17* data is the same as in Figure 2H. G. Violin plot showing expression levels of the example epithelial and fibroblast genes in the CT, the blastema, and the AEC clusters of the Visium dataset. H. Schematics illustrating the AER transcriptional program in developing (axolotl, frog, human, mouse, chicken) (left) and regenerating (axolotl, frog tadpole) limbs (right) of analyzed animals. Red to orange colors is used to indicate changes in the AER transcriptional program, relevant explicitly in the axolotl regeneration schematics. Dash rectangle indicates the zoom-in view of the axolotl regenerating limb with selected AER genes.

Remarkably, our approach identified AEC cells and a subset of axolotl CT cells to have high enrichment scores for the differentially expressed genes in the AER cluster **(Figure S15B)** and gather together in the axolotl regeneration datasets **(Figures 3C and S16-S20)**. Subsequently, we found that the shared genes between these grouped CT cells and AEC includes certain epithelial AER genes (e.g., *Jag2, Cdh1*), although some of the previously reported genes, such as *Fgf8*^7^, were not detected in this population **(Figure 3D)**, which might be due to the limitations to the analyzed dataset (e.g. timing of sample collection, enrichment of non-anterior mesodermal lineage cells). Moreover, we identified these CT cells express AER-associated signaling ligands (e.g., *Bambi, Bmp2*) **(Figures 3D and S21A)**, as well as other genes such as *Vwde, Mdk,* and *Krt18* **(Figure S21B)** that have been already demonstrated to be critical for successful salamander limb regeneration and blastema proliferation^12, 22, 34^.

Then, using our spatial transcriptomics data, we confirmed the presence of the CT cells showing part of the AER program and revealed that they are mostly present in the blastema, where they express both fibroblast- and epithelial-associated genes **(Figures 3E-2G)**, which was also in alignment with the reported mesodermal *Krt5*, *Krt17*, and *Krt18* expression^22, 35, 36^. Moreover, we found that some CT cells showing the AER program are also present in the intact axolotl limbs before limb amputations **(Figure S21C)**. Importantly, in limb development datasets of species analyzed, AER and limb bud mesoderm cells were largely separated **(Figures S16-S20)** and mesodermal cells did not show any enrichment scores for AER-related gene sets **(Figure S15)**. Similarly, we failed to detect the AER program in mesodermal cells during frog tadpole limb regeneration **(Figures S15-S20).** In sum, our analyses revealed an axolotl-specific feature where a part of the AER transcriptional program is also present in the mesodermal lineage cells, which reveals distinctive cellular transcriptional programs for limb development, regeneration, and morphogenesis in general.

## Discussion

The AER is central to successful limb development. However, its presence in salamanders that are widely used to study limb regeneration remained controversial. Advancing from single-gene investigations, our work uses comprehensive cross-species comparisons and clarifies ambiguous propositions related to this topic revealing axolotls have cells with AER transcriptional programs. Nevertheless, contrary to the commonly-accepted assumptions, we found the axolotl AER is not entirely re-formed during limb regeneration to act as the AEC. Given the multiple functional assays demonstrating the essential role of the salamander AEC^12, 37–43^, our results imply that the distinctive axolotl AEC signaling profile might be sufficient for regeneration. Alternatively, genes, other than the ones commonly associated with well-studied signaling pathways, might be critical for the function of the AEC.

Limb regeneration has long been thought to mimic some aspects of limb development^3, 17, 44, 45^, offering a roadmap to establish strategies to regrow lost mammalian limbs. Our results here suggests that more complex and species-specific processes exist for both limb development and regeneration, even among regeneration-competent animals. Notably, axolotl limbs contain parts of the AER program both in ectoderm and mesoderm when compared to other analyzed species - including humans - where only ectodermal cells show an AER program **(Figure 3H**), highlighting significant differences for limb morphogenesis across species. Moreover, identifying the epithelial AER program in the axolotl mesoderm demonstrates an uncommon cross-lineage cellular activity, given the previous studies were not able to identify a lineage switch between ectodermal and mesodermal populations^7, 46–48^, and pose new cell-type evolution and mesodermal plasticity scenarios. Finally, our study provides a non-developmental route to impart limb regeneration to mammals. Inducing epithelial AER programs in CT and ectodermal-derived populations at the same time might be an alternative strategy to regrow lost mammalian limbs.

## Methods

### scRNA-Seq data acquisition and preprocessing

Publicly available raw sequencing data, and the expression matrices of axolotl developing limb datasets were downloaded (Supplementary Table 1). The developmental stages of each sample were determined by the original studies including humans (Carnegie Stage 13), mice (embryonic day (E) 9.5, E10.5, E11.5, E12.5), chickens (Hamburger– Hamilton stage 25), frog tadpoles (Nieuwkoop and Faber (NF) stage 50, NF51, NF52 NF54), and axolotls (stage 50, 52, 54).

For humans, mice, chickens, and frogs datasets, CellRanger (v6.1.1.1) was used for preprocessing the raw data. CellRanger *mkref* function was used to build references for each species with corresponding genome sequences and annotation files (Supplementary Table 1). CellRanger count was used to identify valid cell barcodes, align reads, and quantify gene expression. For axolotl regeneration datasets, kallisto|bustools^49^ was used to generate the expression matrix. Cells passed the filtering in both *DropletUtils::emptyDrops* and *DropletUtils::defaultDrops* were retained for further analyses^50, 51^. The default setting was used unless noted. Cells were further filtered based on dataset-specific thresholds of three metrics: the mitochondrial percentage, the number of transcripts, and the number of genes (Supplementary Table 1).

### Gene list curation of the five analyzed species for comparison

Human, mouse, and chicken orthologs were downloaded from BioMart. As the frog (*Xenopus laevis*) is a pseudo-tetraploid animal, a pseudo-genome was generated: alleles between L and S homologous that are showing the higher expression were considered as the expression^23^. Axolotl genes with the same names as human genes were defined as the orthologs. Multiple axolotl transcripts, thereby genes, could be annotated with the same human gene, for which, only the one with the maximum expression was considered as the expression. In total, 8,855 genes were retained for Figures 1B-1D and S3A and S3B and S14.

### Curation of the gene sets

For cell cycle-related genes, the human gene list was retrieved from a previous study^52^ and orthologs were used for the other species. For signaling ligands, human and mouse genes encoding ligands of FGFs, WNTs, TGFbs, NOTCH/DELTAs, and BMPs signaling pathways were retrieved from CellChat^53^ and orthologs were used for chicken, frog, and axolotl (Supplementary Table 2).

### Clustering of individual scRNA-Seq datasets

Seurat (v4.0.3)^54^ was used for clustering the scRNA-Seq datasets individually. Briefly, the expression matrix was normalized and scaled. The top 2000 highly variable genes were used for principal component analysis (PCA). The first 15 principal components were chosen to build K nearest neighbors (KNN) graph and Louvain clustering. Data were visualized using 2-dimensional UMAP. The default setting was used unless noted. Cell cycle correction was performed when the clustering was biased by the cell cycle (Supplementary Table 1). For this, cell cycle scores were first assigned to each cell using *CellCycleScoring* in Seurat with species-specific cell-cycle genes. Then, the absolute weights of the cell cycle genes (loading values) were summed for each principal component (PC). PCs with values exceeding dataset-specific thresholds were defined as cell cycle-correlated PCs (Supplementary Table 1). PCA was rerun without the top 10% genes from cell cycle-correlated PCs. Specifically, one cluster of low reads count in the chicken HH25 dataset was removed from future analysis.

### Integration of datasets using Seurat

For the multi-species limb atlas, individual datasets were processed into Seurat objects where only one-to-one orthologs of the five species were retained. The annotated cell clusters from each dataset were downsampled to half when exceeding 500 cells in number. The sctransform-based normalization (*SCTransform*) was performed. Genes were ranked by the number of datasets they are deemed variable in (*SelectIntegrationFeatures*) and the top 3000 genes were used to integrate all the datasets (*FindIntegrationAnchors* and *IntegrateData*). Clustering was corrected for the cell cycle effect as described using 1.5 as the threshold to identify cell cycle-correlated PCs. Annotation was performed based on cell-type specific marker genes expression (Figure S3C). Specifically, one confounding cluster spreading across the whole UMAP was removed. The remaining cells were used to determine the final KNN graph (k=30) and UMAP with the top 30 PCs.

For the axolotl regeneration dataset, individual datasets of different time points were integrated into a Seurat object as described without cell downsampling (Figure S11).

### Integration of datasets using SAMap

SAMap (version 1.0.12) was used^55^. The same cells used for Seurat integration were used for SAMap integration. Briefly, pairwise tblastx (version 2.9.0) was performed for transcriptomes of five species using the provided map_genes.sh. BLAST bit scores were used as the initial gene-gene weights. Then, datasets from the same species were concatenated as one input of *samap.run* to perform iterative clustering. In each round of clustering, the gene-gene weights were updated as expression correlations of the matched cells until the alignment score was above the default threshold. The default setting was used unless noted. Cell clusters were defined using the Louvain algorithm implemented in SCANPY^56^ with 2.0 resolution.

### Calculation of integration accuracy

To calculate the integration accuracy, cell type annotations derived from Seurat and SAMap integrated atlases were compared to the annotation of individual dataset clustering results. For this, confusion matrices were generated, using *confusion_matrix* in the cvms R package (1.3.8). The confusion matrices were normalized to the maximum cell number of each cell cluster in the integrated atlases to normalize sample sizes.

### Cluster to lineage and cell type annotation

The lineage or cellular identities of Louvain-defined clusters were defined based on the expression of literature-supported markers (Figures S3C, S10C, S11D, S13E). Ambiguous clusters were labeled as “Unknown”. Specifically in the axolotl regeneration dataset (Figure S11B), the AEC population was determined based on the annotation from the basal ectoderm subclustering (Figures 2B and 2C) where cells from regeneration datasets that fell into the AEC/AEC cluster were defined as AEC. Additionally, the differentially expressed genes (DEGs) of each cell cluster were identified using *FindMarkers* with the default setting and were visualized in Figures S3C, S10C, S11D and S13E.

### Transcriptome-wide comparison of cluster similarity across species

MetaNeighbor was used to calculate the cluster similarity^57^. MetaNeighbor scores cluster similarity with AUROC (area under the receiver operating characteristic). The range of the AUROC scores is from zero to one, where zero indicates dissimilar and one indicates similar. Of note, a value of 0.5 means the algorithm is not able to decide the similarity. The 303 variable genes identified by *MetaNeighbor::variableGenes*, the top 3000 variable genes identified by *Seurat::FindVariableFeatures*, and the top 50 PCs from *Seurat::RunPCA* in the multi-species limb atlas (Figure 1B) were used as input in unsupervised mode, respectively (Figures 1F and S6).

### Single cell gene set enrichment analysis (scGSEA) of the signaling ligands

AUCell^58^ was used for scGSEA. Signaling ligands for each species and the raw count expression of each developing limb dataset were used as input (Supplementary Table 2). AUCell scores were averaged for each species and developmental stage for visualization in Figures 1G and S5A. The average scores were further normalized to the maximum of each dataset and shown in Figure SB.

### Identification of axolotl AER marker gene expression

In the axolotl developing limbs datasets (Figure S2E), the axolotl AER cells were compared to all the other cells and to only the ectodermal cells using *FindMarkers* with the default setting. Axolotl AER cells upregulated genes in all comparisons were intersected, resulting in 14 shared genes (Figure S9A). By visually checking the expression in each cluster, *Dr999-Pmt21178* and *Vwa2* were determined as the most specific ones out of the shared genes (Figure S9B).

### Hybridization chain reaction (HCR) on whole-limb samples

Axolotls were collected during different developmental stages (stages 46, 50 and 53) and fixed with 4% formaldehyde in 1×PBS for 40-60 min and stored in 100% ethanol at −20°C. Fixation was carried out on a rotator at room temperature. Limbs were dissected and HCR protocol was applied as described previously^59^ with modifications. Briefly, limbs were transferred in a new Eppendorf tube containing 500 μl of wash buffer (Molecular Instruments) that has been incubated for 10 min at 37°C. The supernatant was removed and replaced by 500 μl pre-heated hybridization buffer (Molecular probes) for a 30 min incubation at 37°C. During this incubation, the probe solution was prepared by diluting mRNAs targeting probes to 40 nM in 300 μl hybridization buffer and incubated for 30 min at 37°C. Probes for *Vwa2*, and *Dr999-Pmt21178*, designed based on the transcript sequences obtained from the matched transcriptome used in the data analysis (Supplementary Table 1), were purchased from Molecular instruments. The hybridization buffer from samples was taken out and probe solution was placed on samples for a 12 h incubation at 37°C. The samples were then washed twice for 30 min with wash buffer and twice for 20 min with 5×SSC-T on a rotator at room temperature. To visualize probes, amplification solution was prepared by first heating the fluorophore attached hairpins pairs (Molecular instruments) that match to the probes to 95°C for 90 s. Hairpins were then left in the dark at room temperature for 30 min. Afterwards, the final amplification solution was prepared at 72 nM h1 and h2 in 250 μl amplification buffer. Samples were first incubated in amplification buffer without hairpins for 10 min, then placed in the final amplification solution at room temperature, protected from light, for 12-16 h on a rotator. Samples were washed with 2×20 min SSC-T and incubated in 20 μM Hoechst (Sigma, 2261) diluted in 1×PBS at room temperature in dark for 30 min. Finally, the samples were washed 3x10min with PBS and mounted as described below.

### Whole-mount HCR samples imaging

Whole limbs were mounted in 0.8% ultra-low gelling temperature agar (Sigma, A5030) in 1×PBS. Imaging was performed using Confocal imaging was performed using Leica SP8 upright confocal microscope with a 20×/0.75 HC PL Apo air objective and post-processing was performed using ImageJ software.

### Subclustering of the basal ectoderm from axolotl developing and regenerating limbs

Basal ectodermal cell clusters, including the AER clusters, from both axolotl developing limbs (Figure S2E) and regenerating limbs (Figure S11B) were extracted and integrated as one Seurat object as described. Cell cycle correction was performed using 1 as the threshold to define cell cycle-correlated PCs. After removing one cluster with connective tissue markers (*Prrx1*), the KNN graph and the UMAP analysis were re-run using the top 30 PCs (Figure 2B).

### Comparisons between AER and AEC cell clusters

In Figure 2E and 2E, the two-sided Wilcoxon rank sum test was performed on AER (n = 157) and AEC (n = 175) cells. *P* values were corrected with the Benjamini-Hochberg method. Genes with P-values less than 0.05 were considered as significant. Further, DEGs of AER and AEC were calculated using *FindMarkers* function with default parameters (Supplementary Table 3). functional enrichment analysis was performed on the top 200 DEGs (ordered by fold change) of each cell population via Metascape^60^ (Figure S12B).

### Animal Husbandry

Husbandry and experimental procedures were performed according to the Animal Ethics Committee of the State of Saxony, Germany. Animals used were selected by their snout-to-tail sizes.

Axolotl husbandry was performed in the CRTD axolotl facility using methodology adapted from Khattak et al.^61^ and according to the European Directive 2010/63/EU, Annex III, Table 9.1. Axolotls were kept in 18-19°C water in a 12 h light/12 h dark cycle and a room temperature of 20-22°C. Animals were housed in individual tanks categorized by a water surface (WS) area and a minimum water height (MWH). Axolotls of a size up to 5 cm SV were maintained in tanks with a WS of 180 cm2 and MWH of 4.5 cm. Axolotls up to 9 cm SV were maintained in tanks with a WS of 448 cm2 and MWH of 8 cm.

### Spatial transcriptomics

Spatial transcriptomics was performed using the Visium Spatial Gene Expression System (10× Genomics, Pleasanton, CA). Animals 8-9 cm snout to tail were amputated at the level of the lower arm and allowed to regenerate until 5dpa. Limbs were then harvested at the level of the upper arm, fresh frozen in OCT, and then stored at −80°C. Samples were sectioned at −20°C at a thickness of 10µm. Optimization and gene expression assays were carried out according to the manufacturer’s protocol. Briefly, slides were fixed in −20°C methanol, dried with isopropanol, and stained with H&E. A tile scan of all capture areas was generated using an Olympus OVK automated slide scanner system with a color camera and fluorescent module.

For tissue optimization, enzymatic permeabilization was conducted for 0–30 min, followed by first-strand cDNA synthesis with fluorescent nucleotides. The slide was reimaged using the standard Cy3 filter cube. An optimal permeabilization time of 20 min was determined by visual inspection to maximize mRNA recovery while at the same time minimizing diffusion. For gene expression, the initial workflow was similar to the optimization procedure. Library preparation, clean-up, and indexing were conducted using standard procedures. Samples were subjected to pair-ended sequencing using an Illumina Multiplexing generating ∼75M reads.

### Spatial transcriptomics data analysis

Gene expression quantification of the Visium dataset was done using kallisto (0.48.0) and bustools (0.41.0) as previously described^62^. Cells were filtered with *DropletUtils::emptyDrops* and *DropletUtils::defaultDrops* as described. SpaceRanger (v2.0.0) was used to match the Visium spots to the tissue slice. Seurat was used for sctransform-based normalization, PCA, and clustering with the top 30 PCs.

### Identification of AER-specific modules

To define AER cell identity-related transcriptional modules, consensus non-negative matrix factorization (cNMF) (v1.4) was used^33^. The normalized expression matrix with the connective tissue and AER cells from the multi-species limb atlas was used as the input. cNMF requires manually defining the number of modules (controlled by parameter k). We tested k values from 5 to 17, among which k=13 gave the lowest error and decent stability (Figure S14B). Modules 11, 12, and 13 were scored specifically high in only AER cells (Figure S14C) and were defined as AER-specific programs and used for further analyses in Figures S16-S18.

### AER programs in mesodermal lineage cells during limb development and regeneration

To examine if the AER programs are present in limb bud mesoderm and CT, we first defined four gene sets representing the AER programs: (1) DEGs in the AER cluster of the multi-species limb atlas (Figure 1B) which were calculated using *FindMarkers* with the default setting. In total, 545 genes with P-values less than 0.05 and were considered significantly upregulated in multi-species AER cluster. (Supplementary Table 4). (2) three cNMF-derived AER-specific modules that were determined as described above (Figure S14 and Supplementary Table 4).

First, scGESA was performed using AUCell as described using the top 200 genes in the AER DEGs list. AUCell scores were visualized in the UMAP for each dataset (Figure S15). Second, clustering with the top 500 genes from the four AER-related gene sets was performed on each developing and regenerating limb datasets. Only the AER and the limb bud mesoderm clusters in development samples, or the AEC and the connective tissue clusters in regeneration samples, were extracted. Then, PCA was performed using each gene set, respectively. The top 30 PCs were used to project the data into the UMAP (Figures S16-S20).

### Gene ontology analysis of the cNMF-derived AER modules

The top 100 genes from each module were used to perform gene ontology analysis using clusterProfile^63^ (Figure S14C). P-values were corrected using the Benjamini-Hochberg method. GO terms with P-values less than 0.01 were considered as the significantly enriched GO terms.

## Data and code availability

Analysis scripts are available at https://github.com/AztekinLab/Axoltol_AER_2023/tree/master. Requests for data and codes should be addressed to C.A..

## Acknowledgments

We thank Aztekin and Sandoval-Guzman Labs for the discussions; M. Gurel and M. Brbic for discussions on factorization and module identification; M. Ros, D. Suter, G. La Manno, and M. Gurel for their critical reading of the manuscript; A. Petzold and the CMCB Genome Facility for help with spatial transcriptomics sequencing and data processing.

## Author contributions

Conceptualization: mainly C.A., J.Z.

Methodology: C.A., J.Z.

Software: J.Z.

Formal analysis: J.Z.

Investigation: C.A., J.Z., G.T., E.S., R.A.

Project administration: C.A.

Supervision: mainly C.A., T.S.G.

Funding acquisition: C.A., T.S.G.

Resources: K.B.

Data curation: C.A., J.Z.

Writing – original draft: mainly C.A., J.Z.

Writing – review and editing: all authors.

---

**Figure S1.**
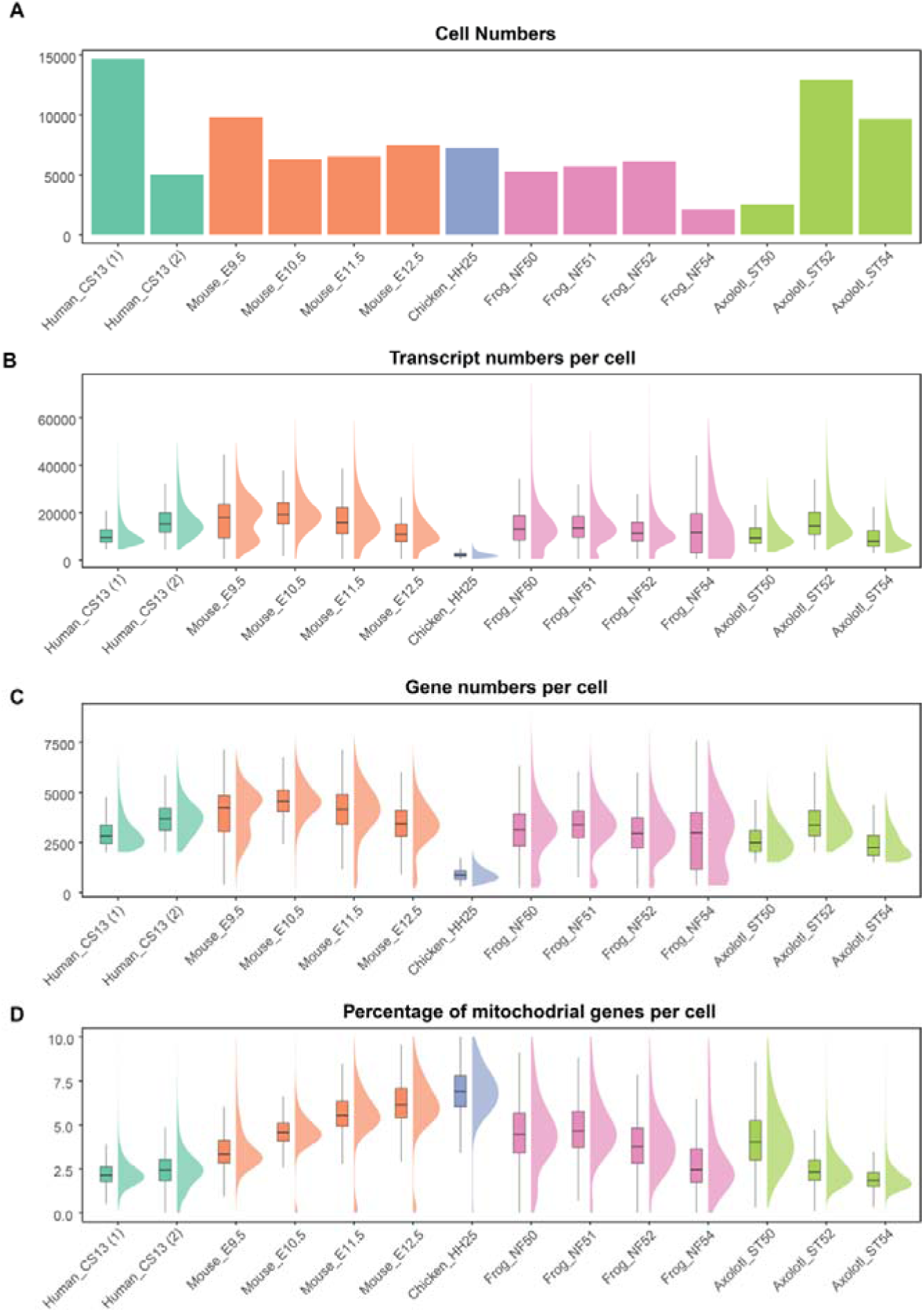
Quality assessment of the re-analyzed publicly available limb development datasets of five species. A. Barplot showing the used cell numbers after filtering in each dataset is visualized. Two human datasets of the same developmental stage from independent studies were used and denoted as Human_CS13 (1) and Human CS13(2). Please see Table 1 for full details. B. Boxplot showing the transcript number per cell in each dataset is visualized. Please see Table 1 for full details. C. Boxplot showing the gene numbers per cell in each dataset is visualized. Please see Table 1 for full details. D. Boxplot showing the percentage of mitochondrial genes per cell in each dataset is visualized. Please see Table S1 for full details.

**Figure S2.**
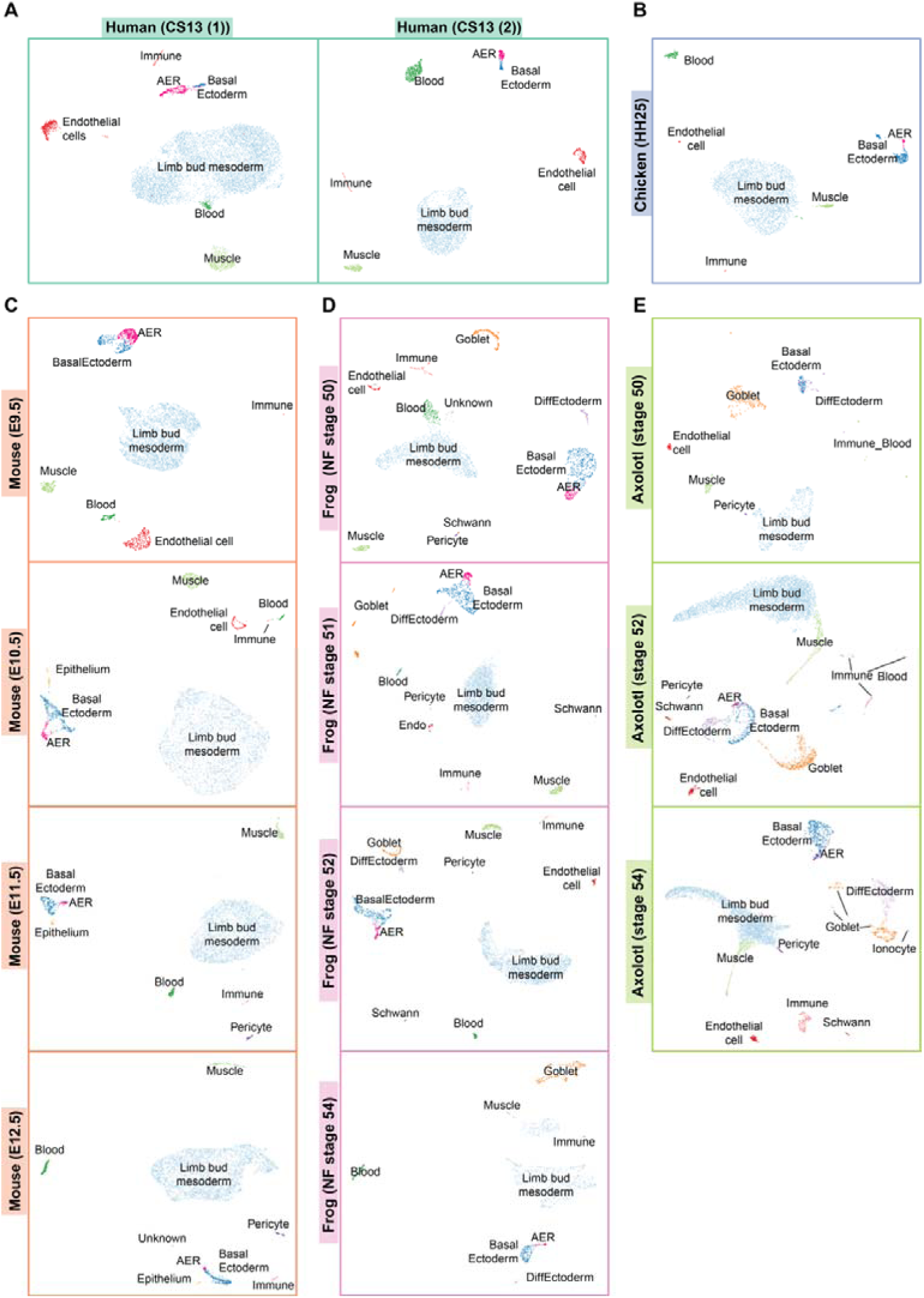
UMAP visualization of the re-analyzed individual datasets of limb development. UMAP visualization of individual limb development datasets for (A) human, (B) chicken, (C) mouse, (D) frog, and (E) axolotl are visualized. Two human datasets of the same developmental stage from independent studies were used and denoted as Human_CS13 (1) and Human CS13(2). Lineage and cell type annotations are based on marker genes in Figure S3, and labeled by different colors and text, except the axolotl AER cells, which are identified based on Figure 1D and re-plotted in this figure.

**Figure S3.**
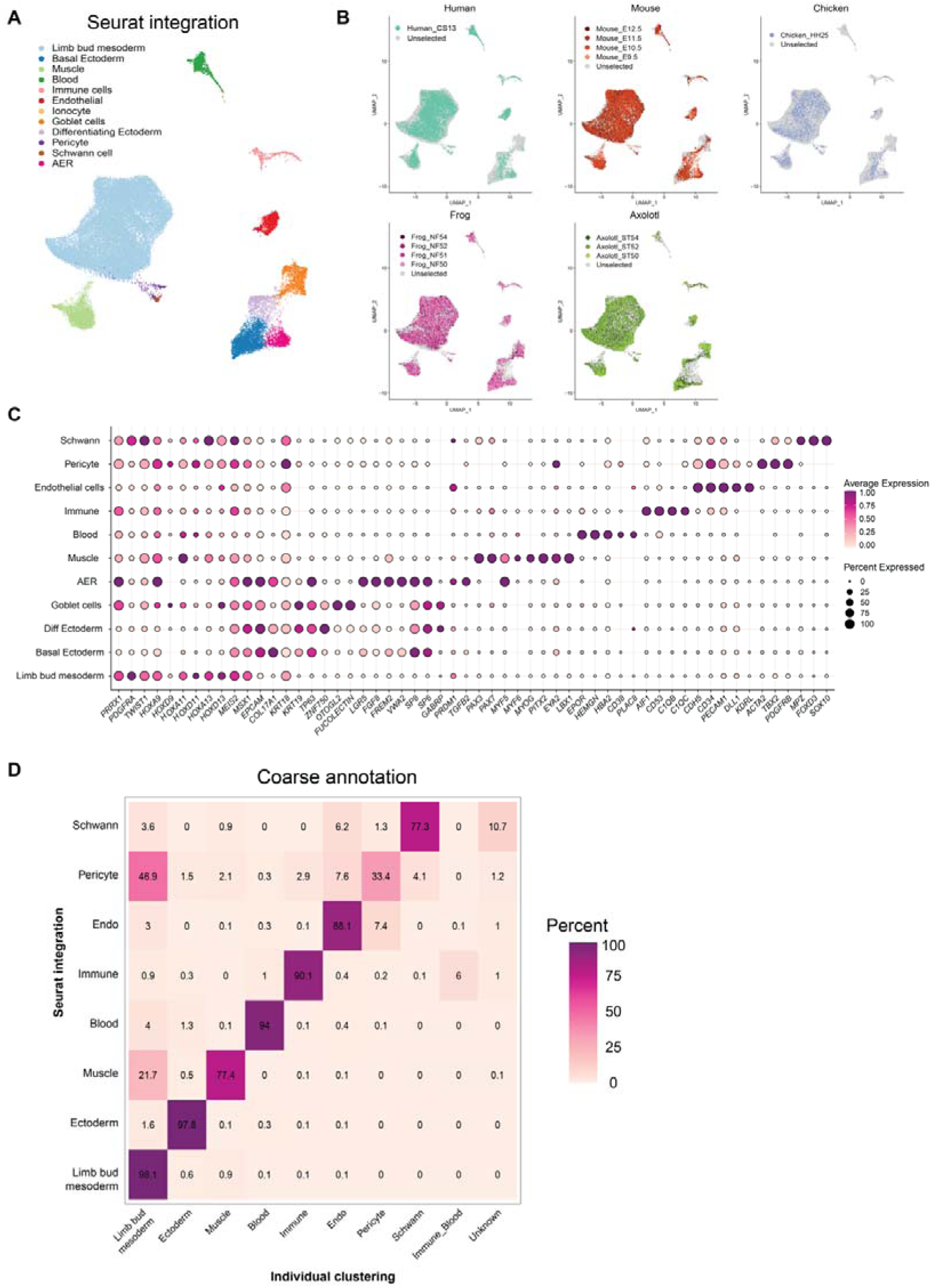
Seurat-integrated multi-species limb atlas. A. UMAP plot of Seurat-integrated multi-species limb atlas is shown. Individual datasets from each species and developmental stage (full list in Figure S2 and Supplementary Table 1) are integrated. Cell lineage and cell type identities are labeled by different colors and text. Please note that this plot is the same as in Figure 1B. B. UMAP plots of species and developmental stage contribution to the Seurat-integrated multi-species limb atlas are shown for each species. Species are color-coded and developmental stages are indicated with a shading of the same color. C. Dotplot showing used marker genes to annotate clusters. The dot color indicates the mean expression that was normalized to the max of each cell type and to the max of each gene; the dot size represents the percentage of cells with non-zero expressions. D. A confusion matrix is plotted to determine the annotation accuracy for cells between the Seurat-integrated multi-species atlas and individual maps from Figure S2.

**Figure S4.**
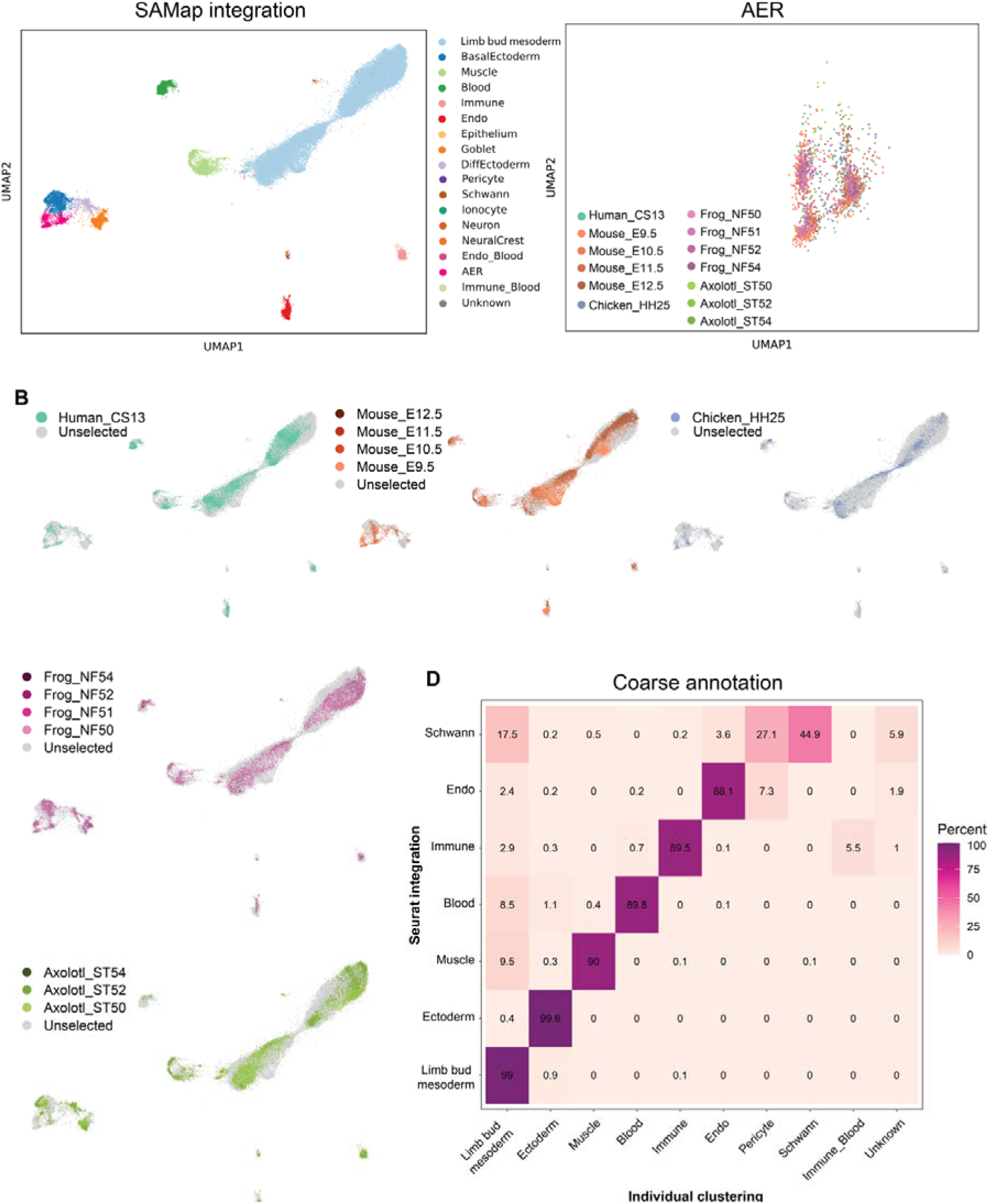
SAMap-integrated multi-species limb atlas. A. UMAP plot of SAMap -integrated multi-species limb atlas is shown. Individual datasets from each species and developmental stage (full list in Figure S2 and Supplementary Table 1) are integrated. Clustering and annotation based on marker gene expressions (please see Figure S3) are indicated lineage and cell type identities, and labeled by different colors and text. B. UMAP plots of species and developmental stage contribution to the SAMap-integrated multi-species limb atlas are shown for each species. Species are color-coded and developmental stages are indicated with a shading of the same color. C. UMAP plot of species contribution to the SAMap-integrated multi-species limb atlas AER cluster is visualized. Cells from different species are color-coded and the developmental stage is indicated. D. A confusion matrix is plotted to determine the annotation accuracy between the SAMap-integrated multi-species atlas and individual maps from Figure S2.

**Figure S5.**
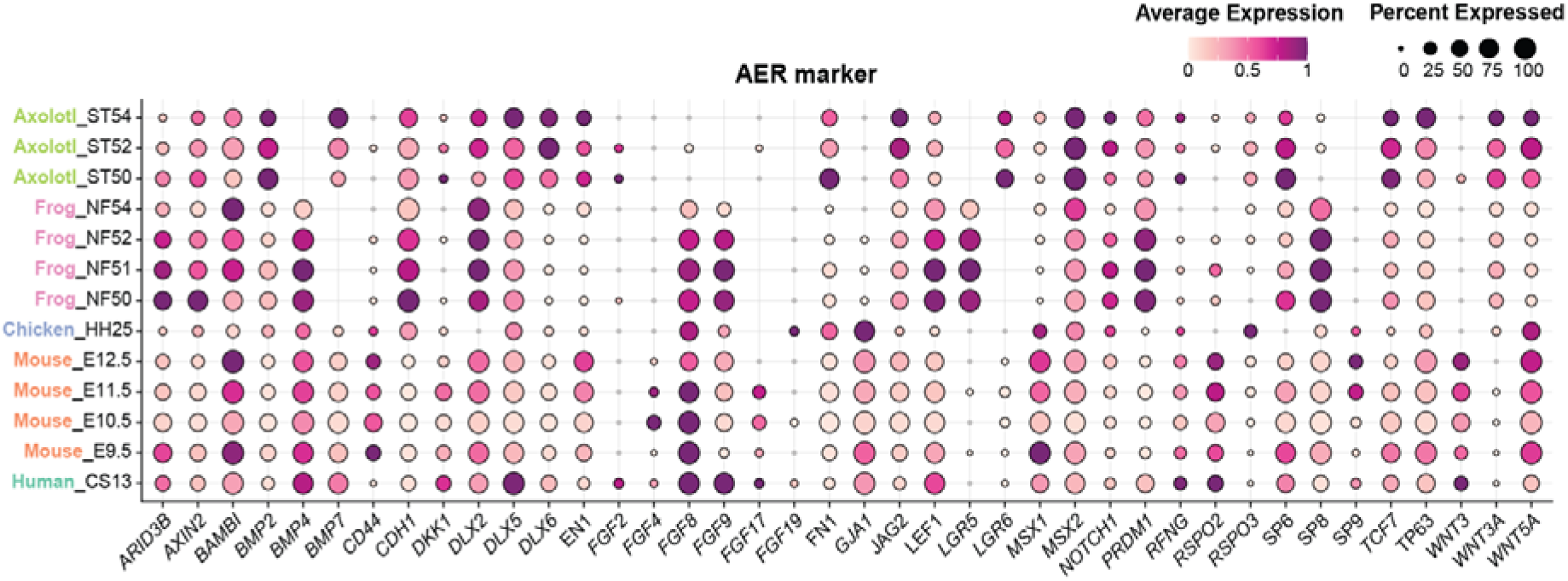
AER marker expressions in AER cells of the analyzed species. Dotplot showing an extended list of AER markers. The dot color indicates the mean expression that was normalized to the max of each cell type and to the max of each gene; the dot size represents the percentage of cells with non-zero expression. Please note that some of the genes in this figure and their values are the same as in Figure 1E.

**Figure S6.**
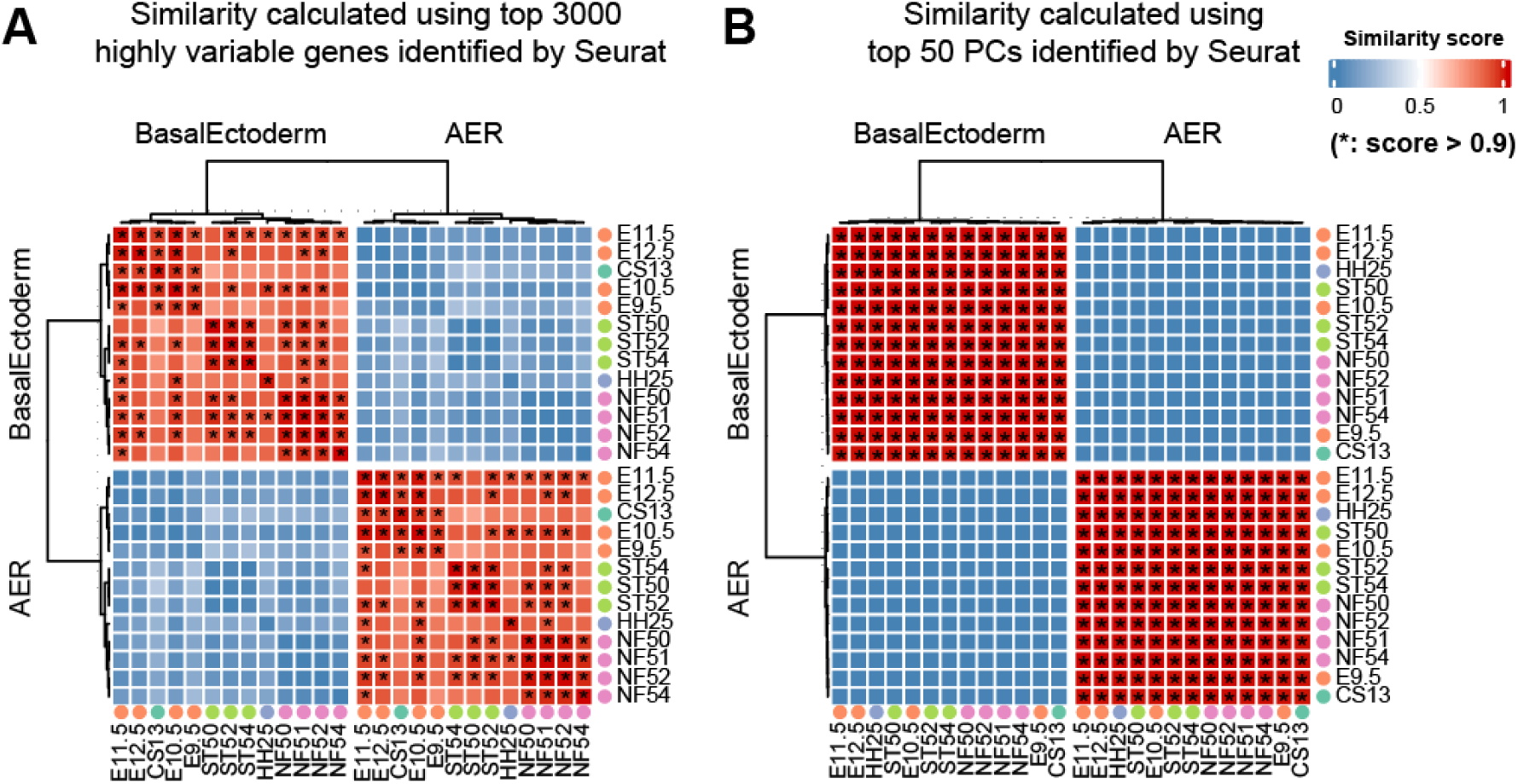
Basal ectoderm and AER clusters similarities of the analyzed species. Heatmaps showing the pair-wise similarity scores of basal ectoderm and AER clusters across species. X and Y axes indicate different species with their developmental stages. Similarity scores are calculated by using (A) the top 3000 highly variable genes, and (B) the top 50 principal components. Asterisks (*) denote the pairs with similarity scores above 0.9. AER clusters are compared to basal ectoderm clusters to highlight specific transcriptome profiles.

**Figure S7.**
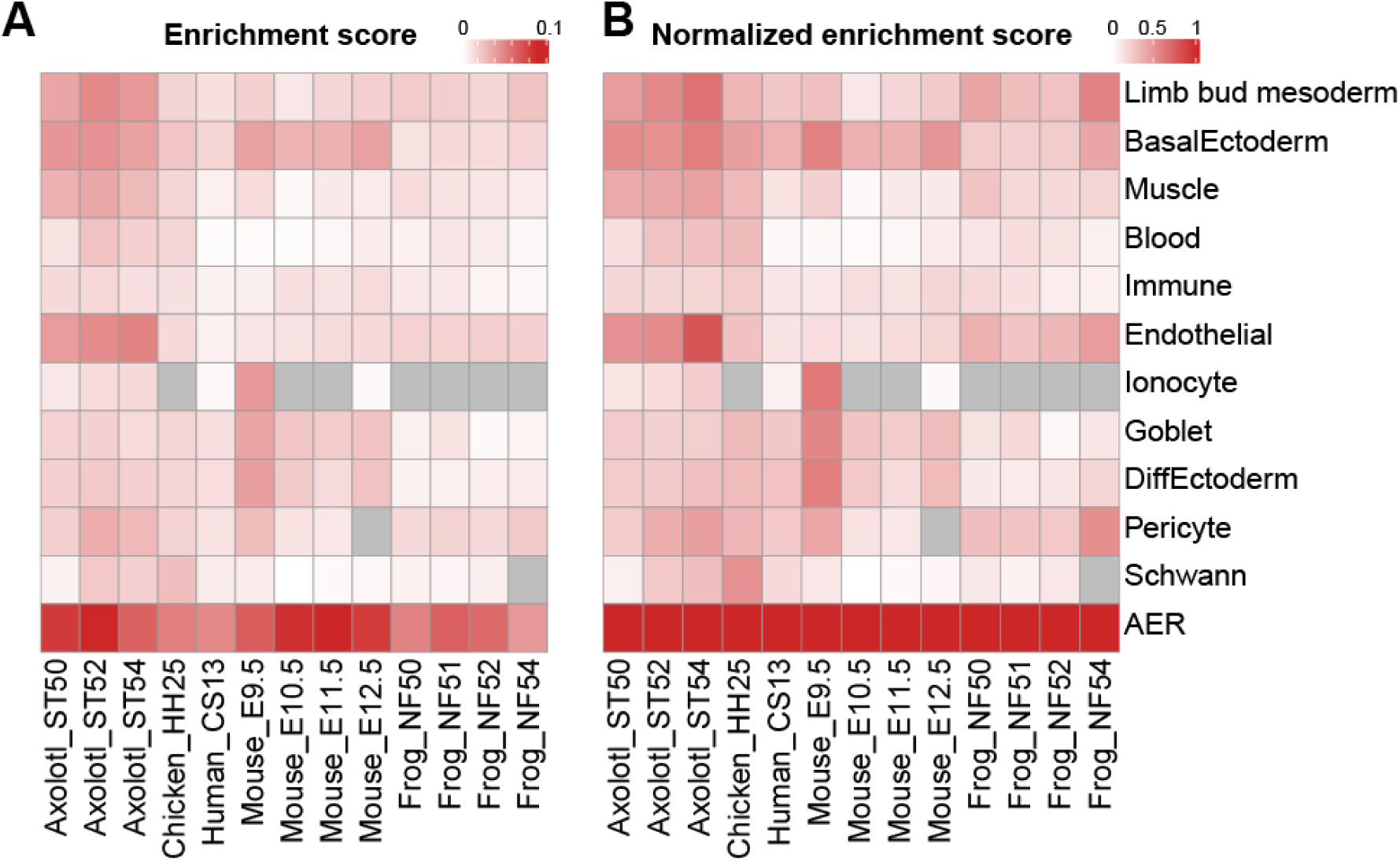
Signaling ligand gene set enrichment scores for lineages and cell types in the analyzed species. Heatmap showing the (A) calculated and (B) max-normalized signaling ligand gene set enrichment scores for each annotated population in the multi-species limb atlas (Figure 1B). Grey indicates missing cell cluster. Please note that the same data for the basal ectoderm and the AER clusters are shown in Figure 1G.

**Figure S8.**
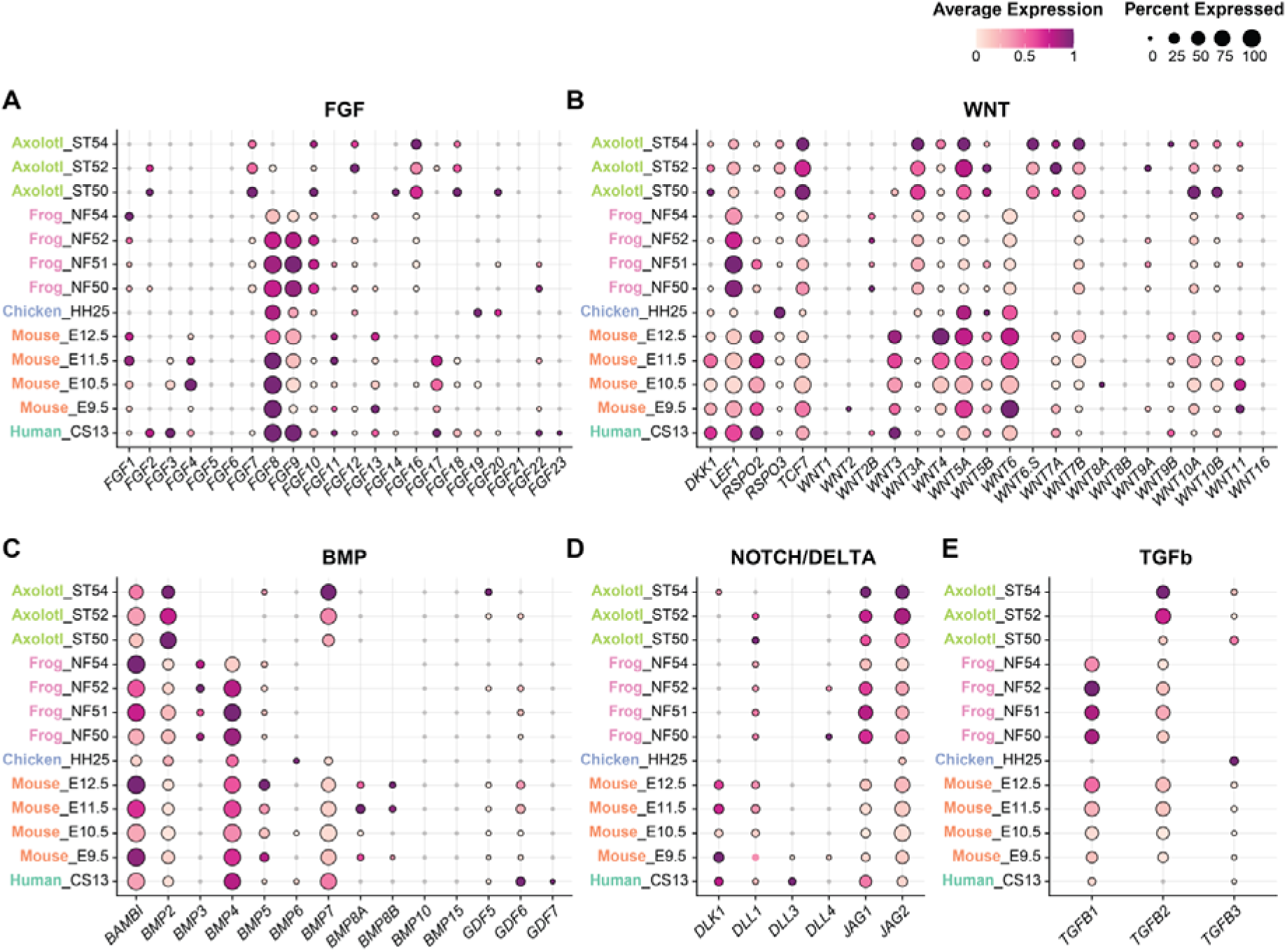
Signaling pathways-associated gene expressions in AER cells in the analyzed species. Dotplots showing expression profiles of various genes from FGF (A), WNT (B), BMP (C), NOTCH (D), and TGFb (E) pathways. The dot color indicates the mean expression that was normalized to the max of each cell type and to the max of each gene; the dot size represents the percentage of cells with non-zero expression.

**Figure S9.**
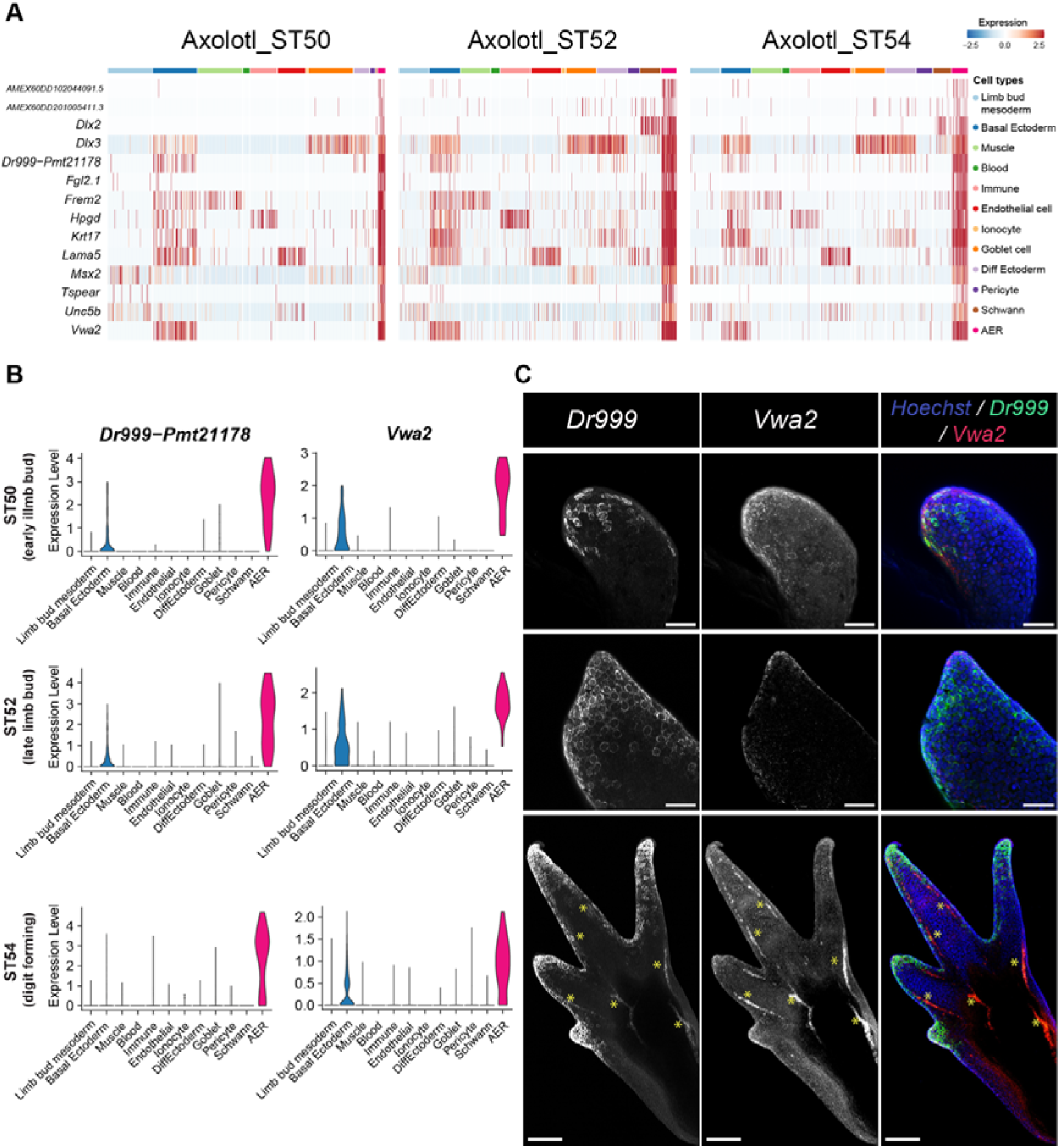
Spatial organization of axolotl AER cells during limb development. A. Heatmap showing the expression of 14 shared AER-specific differentially expressed genes in different cell types in axolotl developing limbs. Stages are indicated on the top. B. Violin plots showing expression levels of the putative axolotl AER markers are shown. Please note that the analyzed scRNA-Seq datasets are for hindlimbs, and morphologies of corresponding developing limbs are indicated in the figure. C. Max-projection confocal images of Stage 46 (top), 50 (mid), and 53 (bottom) axolotl forelimbs correspond to ST50, 52, 54 hindlimb scRNA-Seq data, respectively, stained for *Dr999-Pmt21178* and *Vwa2* mRNA via HCR. In the merged figures green corresponds to *Dr999-Pmt21178* mRNA *(referred to as Dr999 in the figure)*, red corresponds to *Vwa2* mRNA, and blue corresponds to Hoechst. Yellow * is added to the bottom images to indicate autofluorescence. Scale bar for top and mid images: 100 μm; scale bar for bottom images: 200 μm.

**Figure S10.**
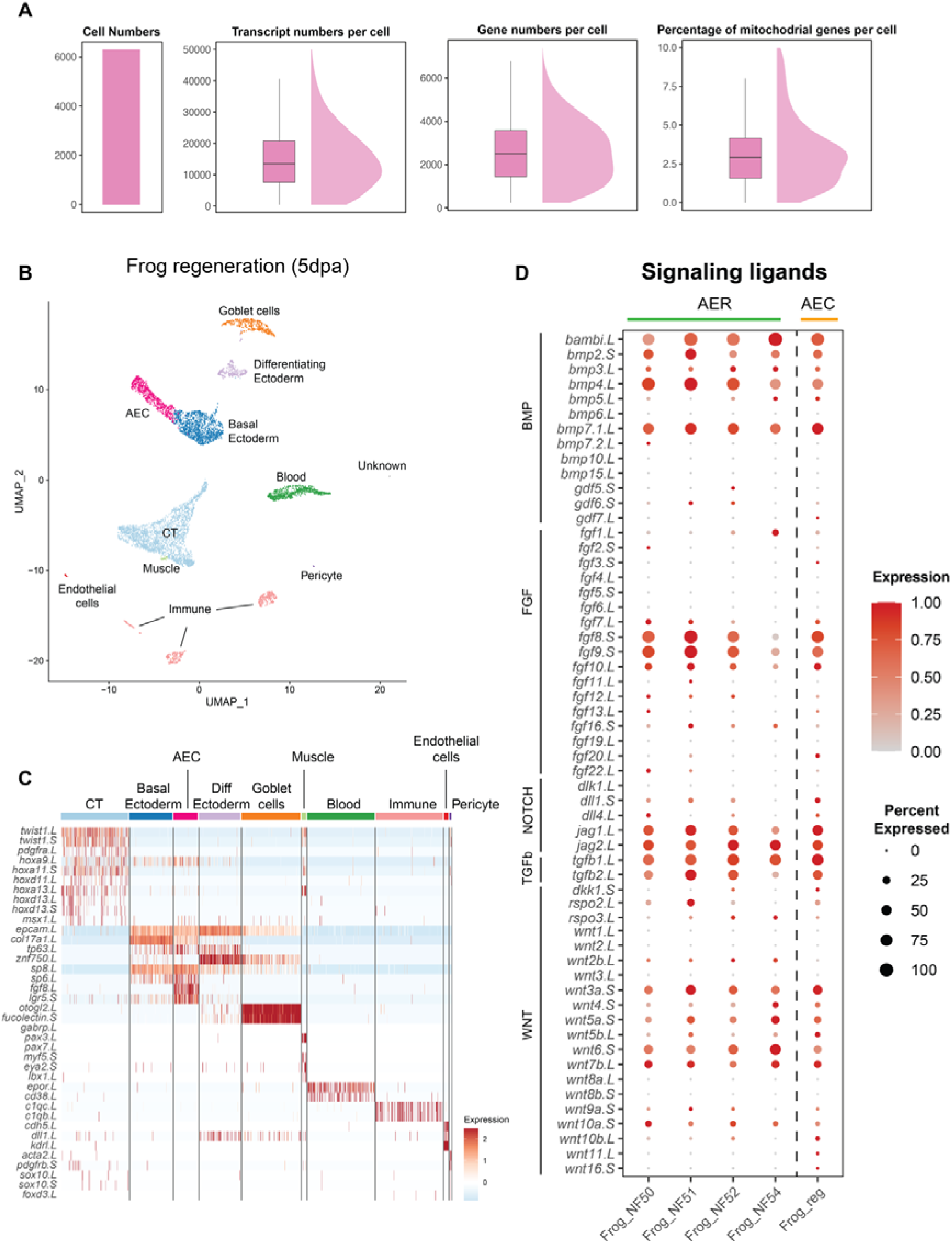
Quality assessment of the re-analyzed publicly available *Xenopus* limb regeneration dataset. A. Barplots showing the used cell numbers, the transcript number per cell, the gene numbers per cell, and the percentage of mitochondrial genes per cell after filtering in *Xenopus laevis* 5 days-post amputation (dpa) limb regeneration dataset are visualized. B. UMAP plot of the identified and annotated clusters. Clusters were annotated based on marker genes listed in Figure S10c. C. Heatmap showing the expression profile of marker genes that are used to annotate clusters. D. Dotplot showing signaling ligand expressions in the AER or AEC clusters during *Xenopus laevis* limb development and regeneration, respectively. Please note that a similar figure was generated in Aztekin et al, 2021, but in this figure, NF Stage 50, 51, and 52 from Lin et al. 2021 were used.

**Figure S11.**
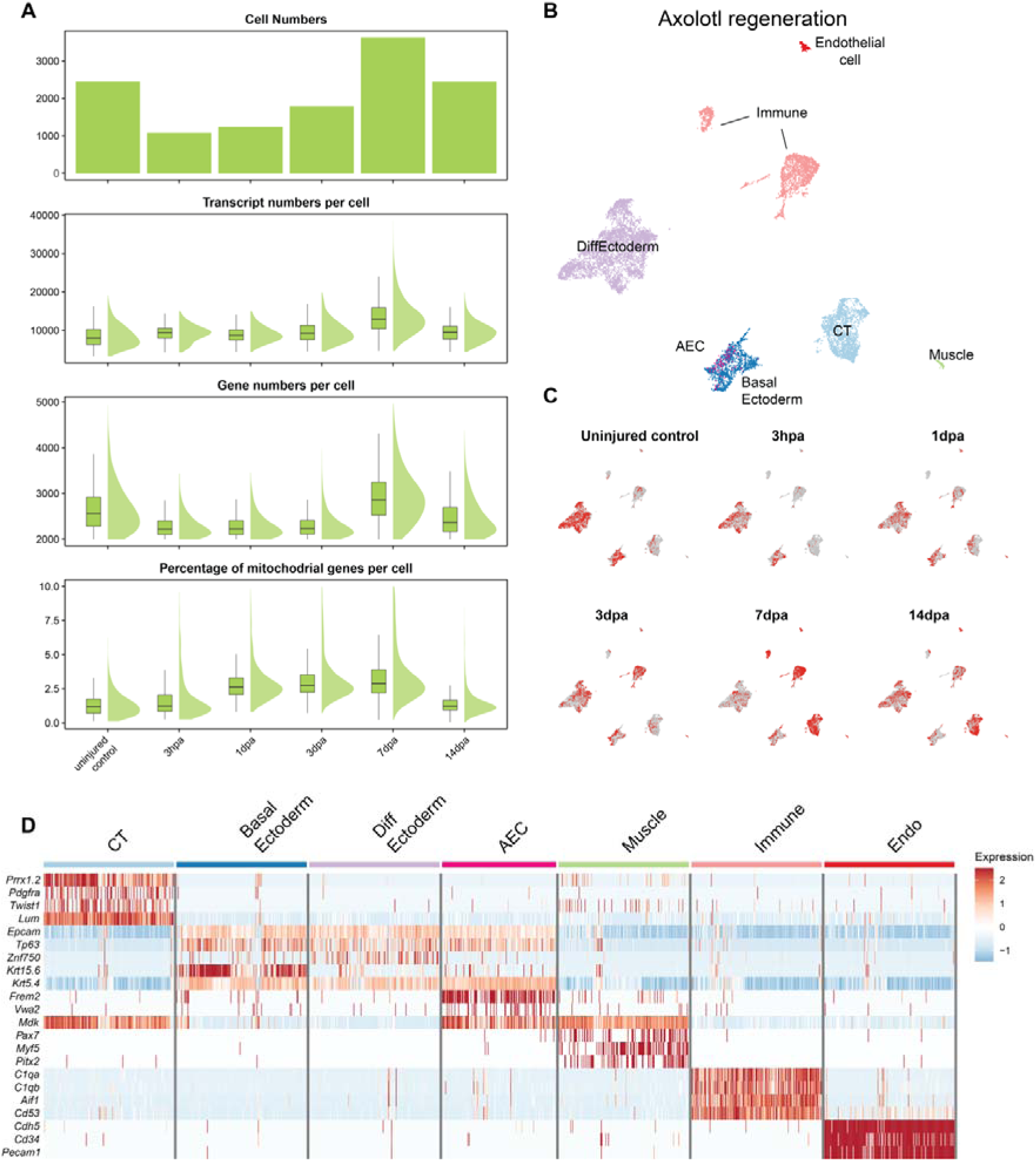
Quality assessment of re-analyzed publicly available axolotl limb regeneration dataset. A. Barplots showing the used cell numbers, the transcript number per cell, the gene numbers per cell, and the percentage of mitochondrial genes per cell after filtering in axolotl limb regeneration datasets are visualized. B. UMAP plot of the identified and annotated clusters. Clusters were annotated based on marker genes listed in Figure S11d. The AEC cells were identified by the analysis described in the text and in Figures 2A-2C, then re-plotted and colored dark pink in this figure. C. Sample contribution to the axolotl regeneration dataset is visualized. Red dots indicate cells from the selected sample; gray dots indicate the other cells. hpa: hours-post amputation; dpa: days-post amputation. D. Heatmap showing the expression profile of marker genes that are used to annotate clusters, except AEC cells and their marker expressions indicated following the analysis indicated in text and Figures 2A-2C.

**Figure S12.**
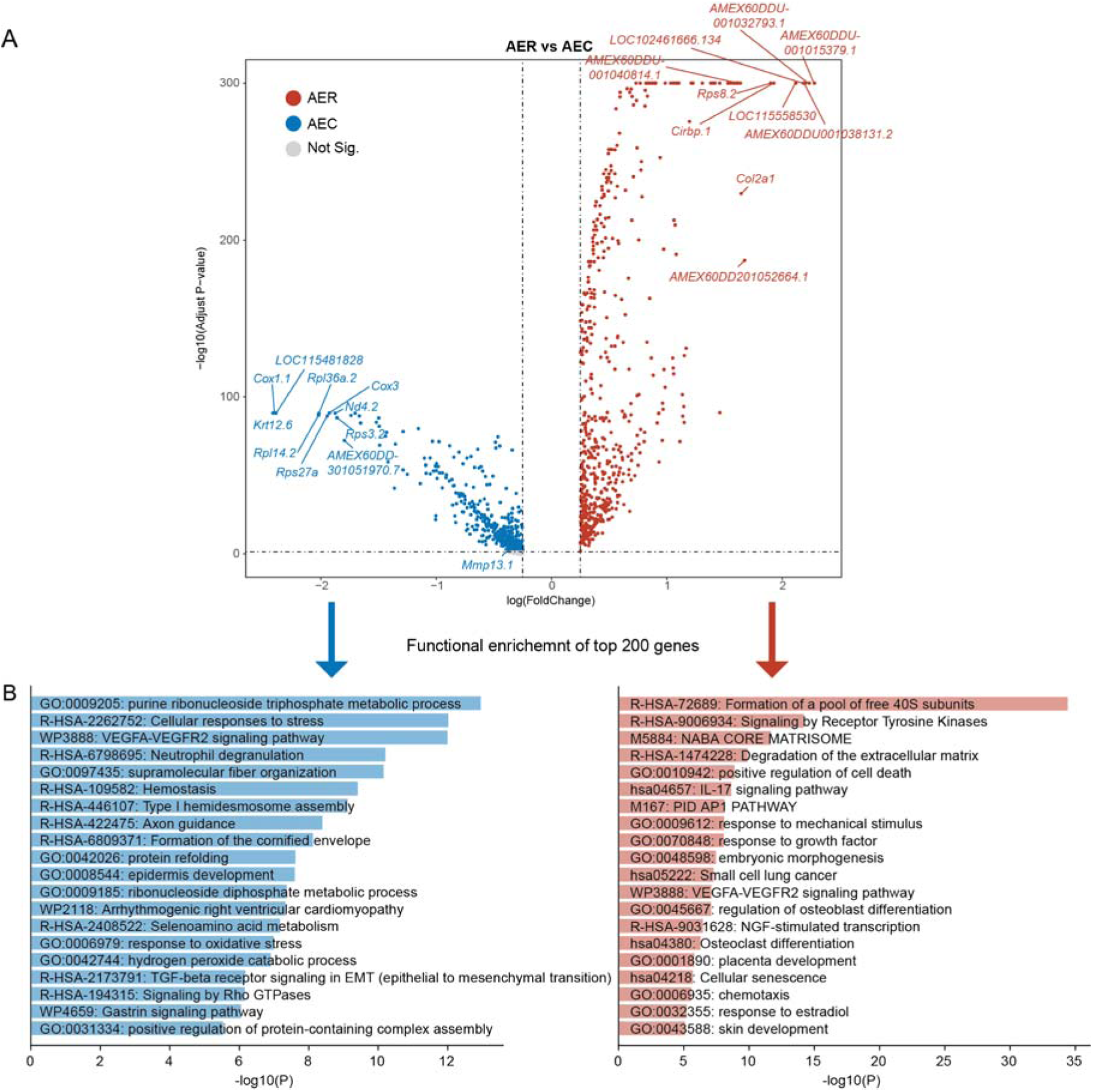
Differentially expressed gene analysis between axolotl AER and AEC cells. A. Volcano plot showing the differentially expressed genes (DEGs) between axolotl AER and AEC populations. Red and blue dots indicate genes significantly enriched in the AER or the AEC, respectively. Gray dots indicate statistically not significant genes. The top 10 DEGs are labeled. B. Barplots showing enriched GO terms based on the top 200 DEGs (ordered by fold change) in AER (red-left) and AEC (blue-right) cells.

**Figure S13.**
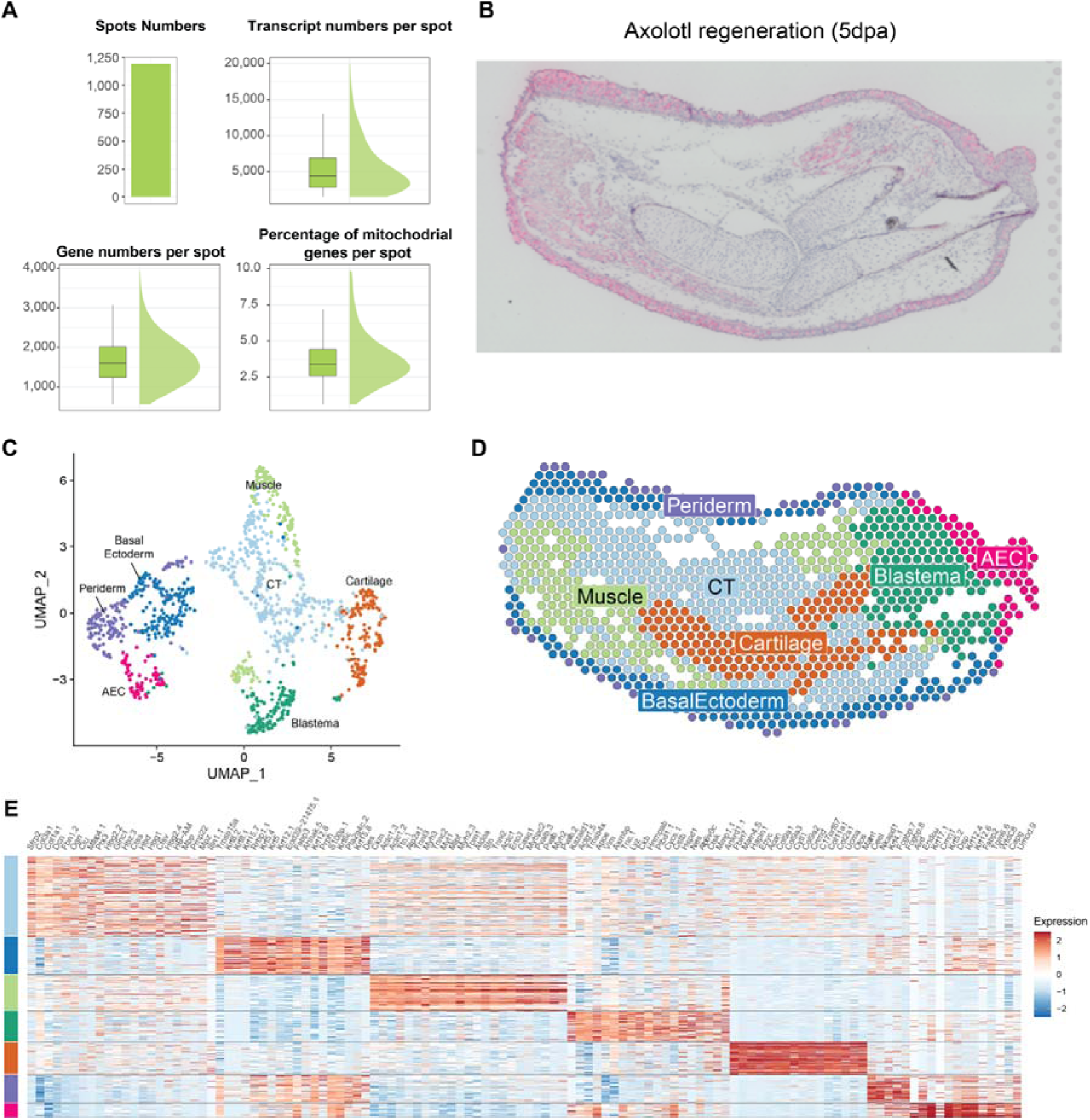
Quality assessment and clustering results of axolotl 5 days-post-amputation 10X Visium spatial transcriptomics (Visium) dataset. A. Barplots showing used Visium spot numbers, transcript number per spot, gene numbers per spot, and the percentage of mitochondrial genes per spot after filtering are visualized. B. 5 dpa axolotl limb regeneration tissue section that is used for the Visium is shown and stained for hematoxylin and eosin. Please note that this image is the same as Figure 2F but added in this figure for easier visual comparison to Figure S13D, which contains full annotation. C. UMAP plot of the Visium clusters is shown. Clusters were annotated by marker gene expressions (Figure S13e) and confirmed by morphologies in hematoxylin and eosin staining on the tissue section in Figures 2F and S13B. The identified clusters are color-coded and tissue annotation is labeled. D. Visium spots were colored by tissue types identified in Figure S13C. E. Heatmap showing the top 20 DEGs (ordered by fold change) for the identified clusters is shown.

**Figure S14.**
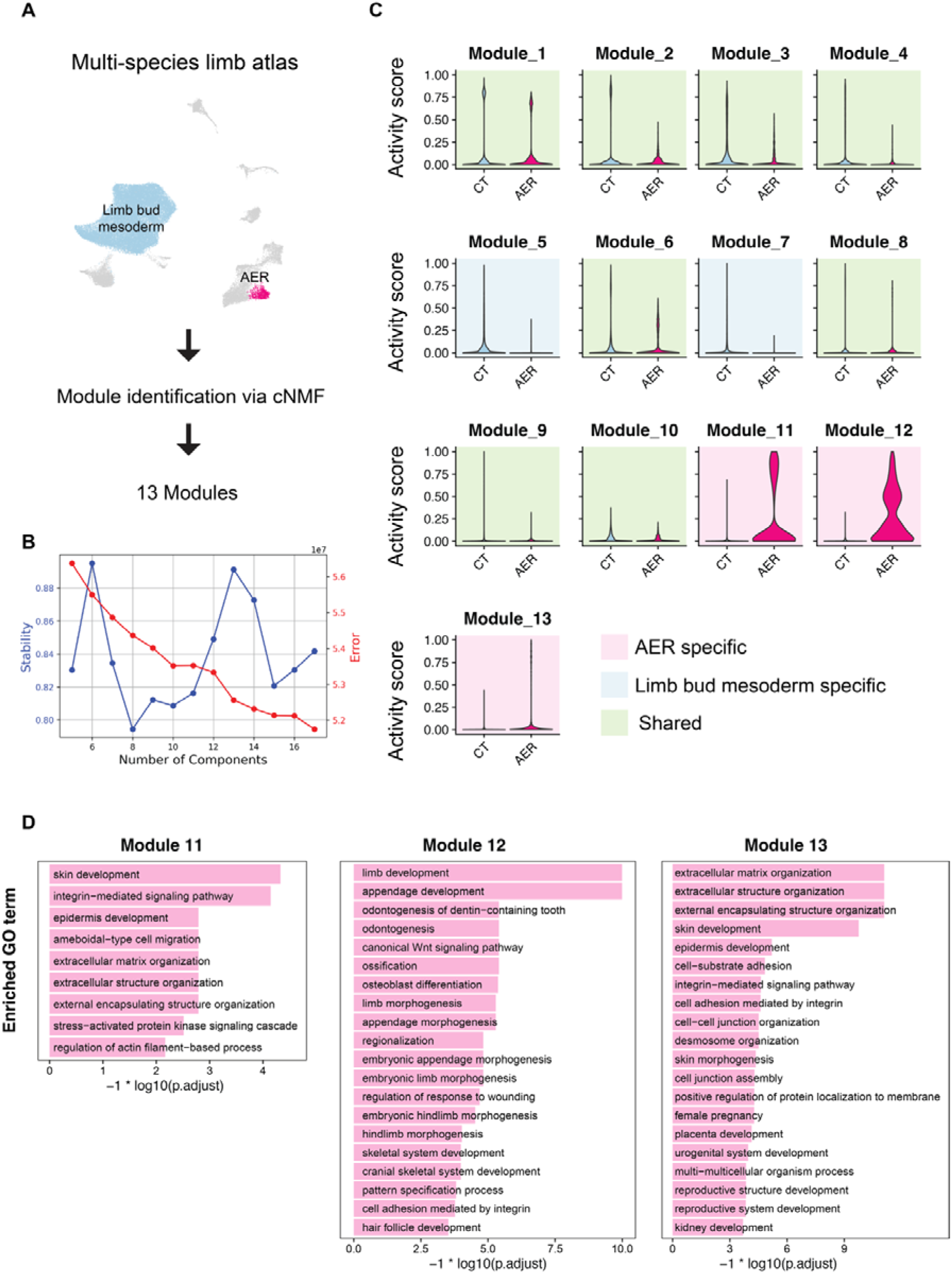
Consensus nonnegative matrix factorization (cNMF) to identify AER-related modules. A. The strategy to perform consensus negative matrix factorization (cNMF) is described. cNMF was used on AER and CT clusters to identify cell identity and cell activity modules. B. Stability (blue line) is measured by the Euclidean distance silhouette score of the clustering and Frobenius error of the consensus solution (red line) for each tested k value (module number) shown in the X axis. Please see Methods for more detail. C. Violin plots showing activity scores for identified cNMF modules in CT and AER clusters. Modules specific to AER, CT, or shared are color-coded: pink, AER-specific modules; blue, CT-specific modules; green, shared modules. D. Enriched GO terms are determined for the top 100 genes (ordered by gene expression program scores) in the identified AER-specific modules. Barplot showing the top GO is visualized, and GO terms are labeled.

**Figure S15.**
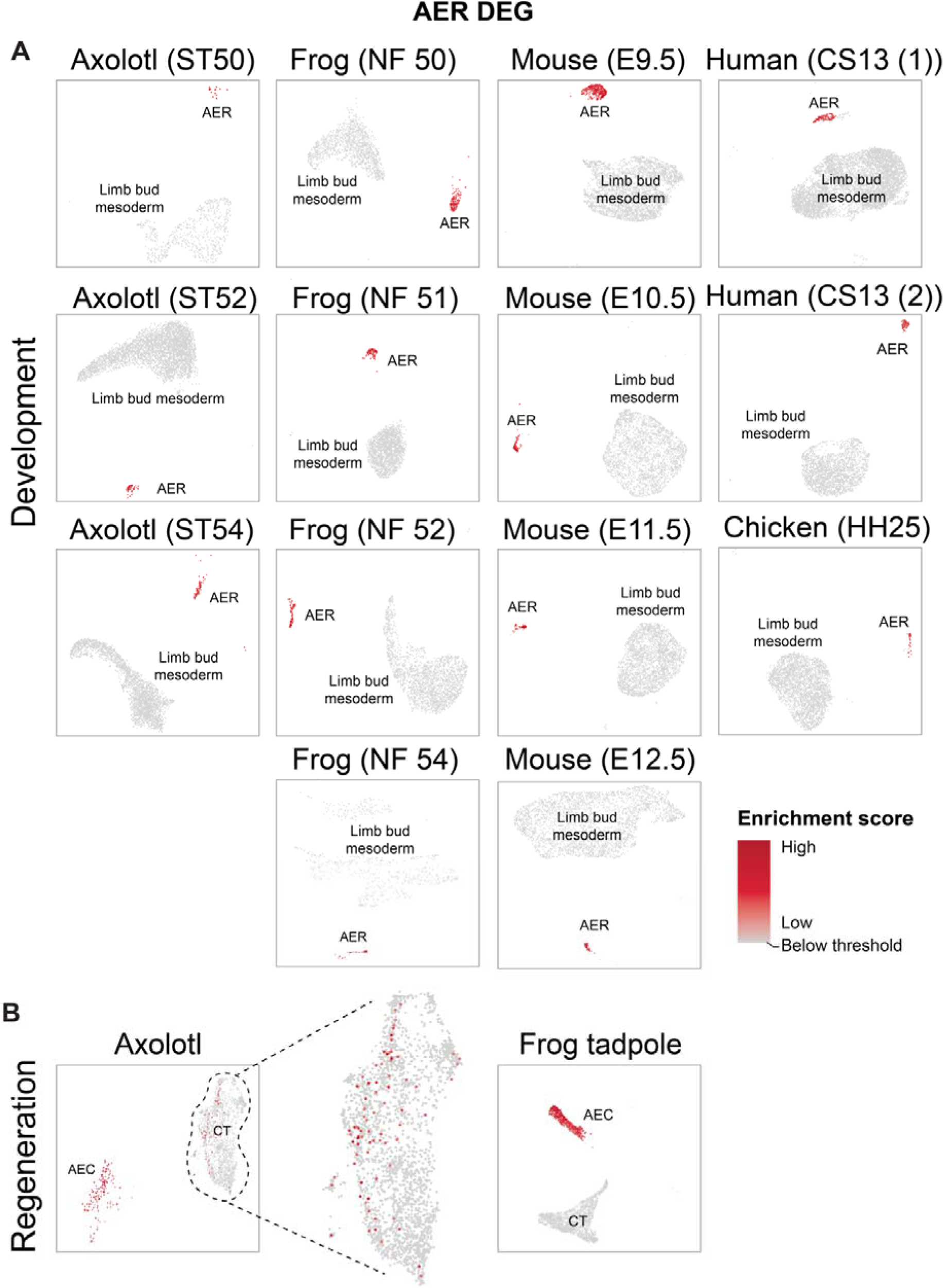
Single cell gene set enrichment analysis (scGSEA) using differentially expressed AER gene sets for individual datasets. UMAP visualization of the scGSEA of indicated datasets of limb development (A) and regeneration (B) based on the top 500 AER DEGs in multi-species limb atlas (Figure 1B). The enrichment scores are indicated in shades of red. Cells failed to pass the enrichment threshold are colored in grey.

**Figure S16.**
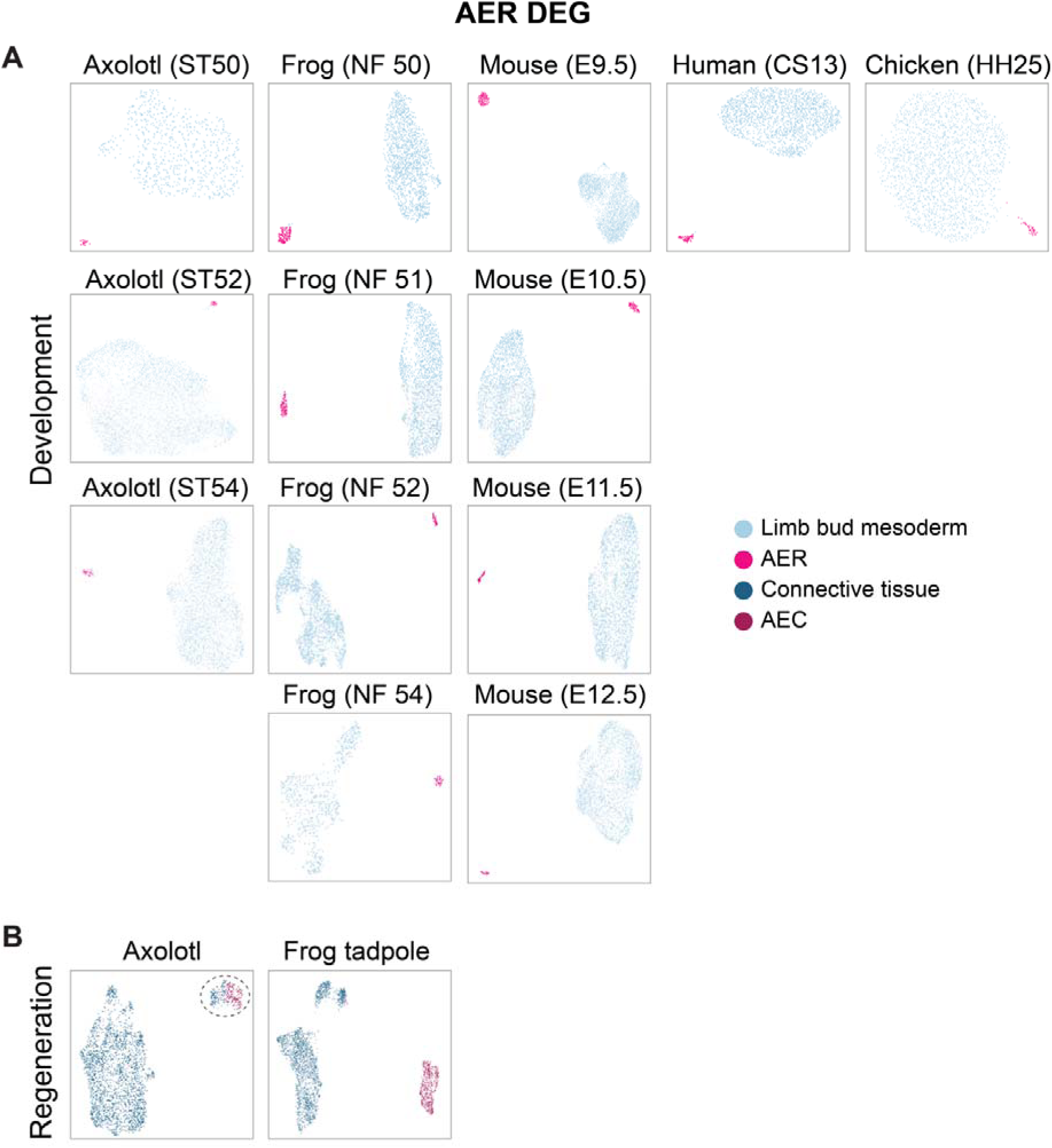
Clustering based on differentially expressed AER gene sets for individual datasets. UMAP visualization of the clustering of indicated datasets of limb development (A) and regeneration (B) based on the top 500 AER DEGs in multi-species limb atlas (Figure 1B) for individual datasets. In development datasets: light blue, limb bud mesoderm cells; pink, AER cells. In regeneration datasets: dark blue, CT cells; dark pink, AEC cells. Connective tissue cells gathered with the AEC population are highlighted with a dashed line. Please note that Mouse E10.5, chicken E4.5, human CS13, frog ST51, and axolotl ST52 development datasets, and the axolotl and frog tadpole regeneration datasets are also shown in Figures 3B and 3C.

**Figure S17.**
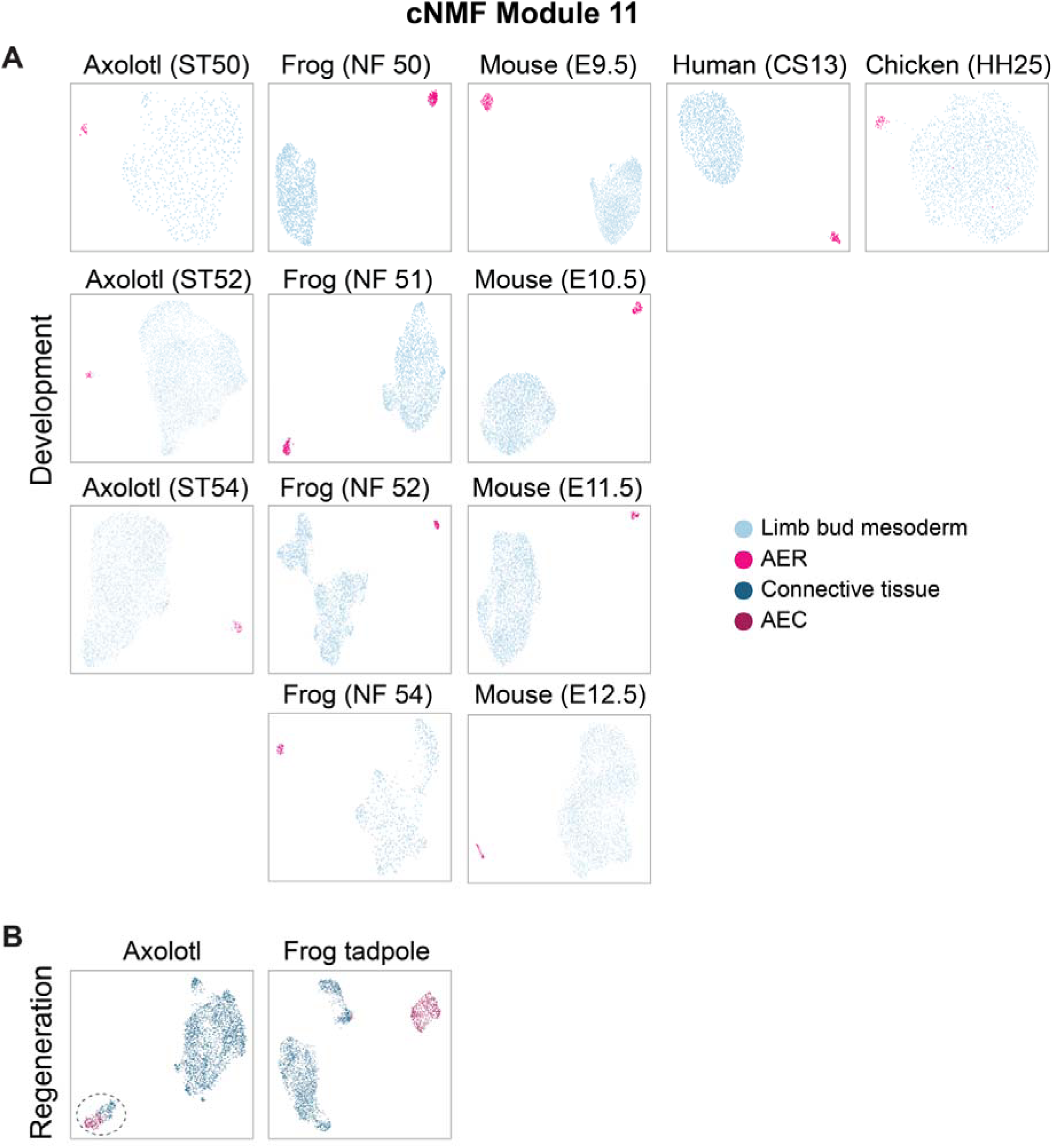
Clustering based on cNMF module 11 for individual datasets. UMAP visualization of the clustering of indicated datasets of limb development (A) and regeneration (B) based on the top 500 genes from AER-specific cNMF module 11 (Figure S14 and Supplementary Table 4). In development datasets: light blue, limb bud mesoderm cells; pink, AER cells. In regeneration datasets: dark blue, CT cells; dark pink, AEC cells. Connective tissue cells gathered with the AEC population are highlighted with a dashed line.

**Figure S18.**
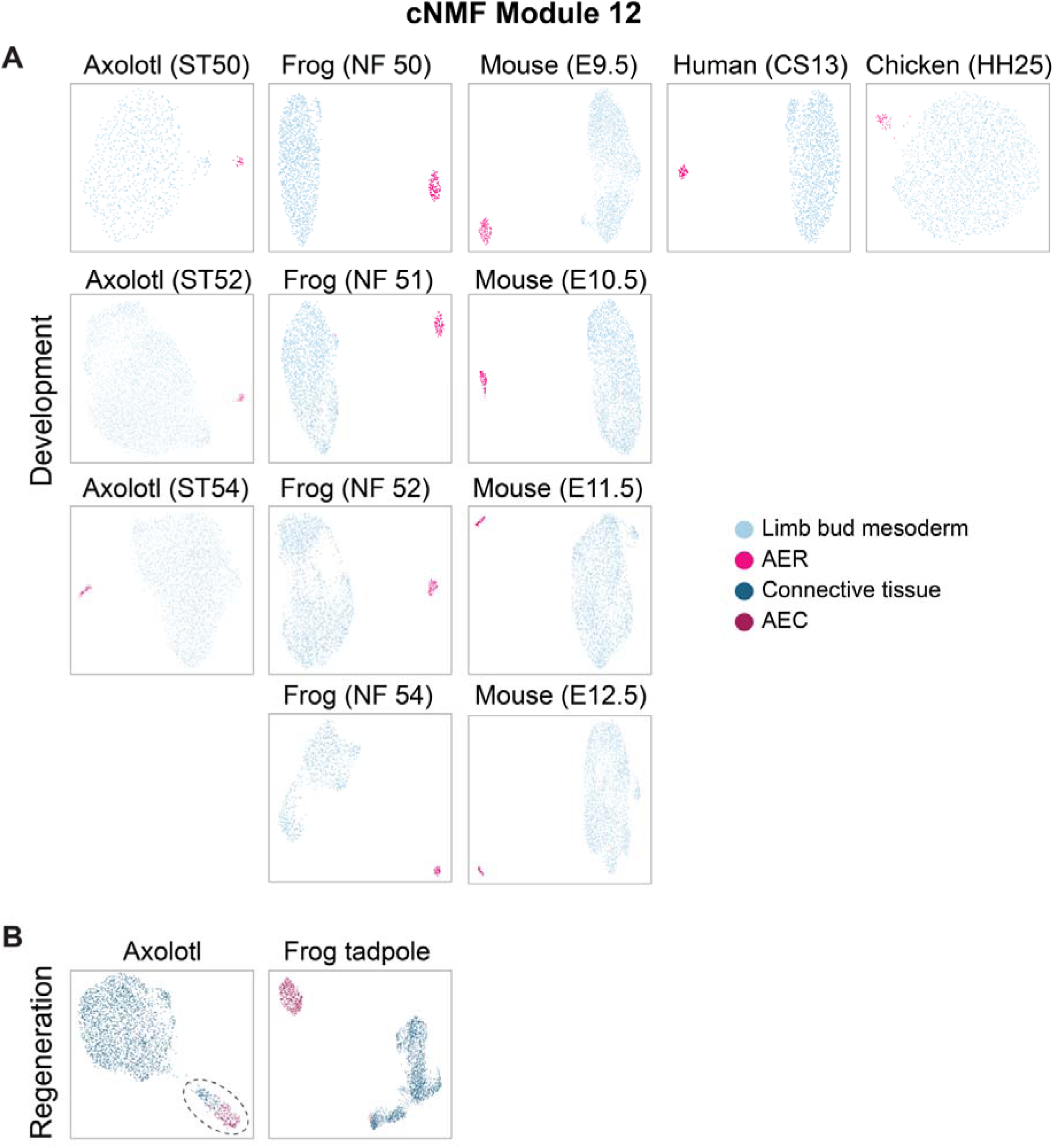
Clustering based on cNMF module 12 for individual datasets. UMAP visualization of the clustering of indicated datasets of limb development (A) and regeneration (B) based on the top 500 genes from AER-specific cNMF module 12 (Figure S14 and Supplementary Table 4). In development datasets: light blue, limb bud mesoderm cells; pink, AER cells. In regeneration datasets: dark blue, CT cells; dark pink, AEC cells. Connective tissue cells gathered with the AEC population are highlighted with a dashed line.

**Figure S19.**
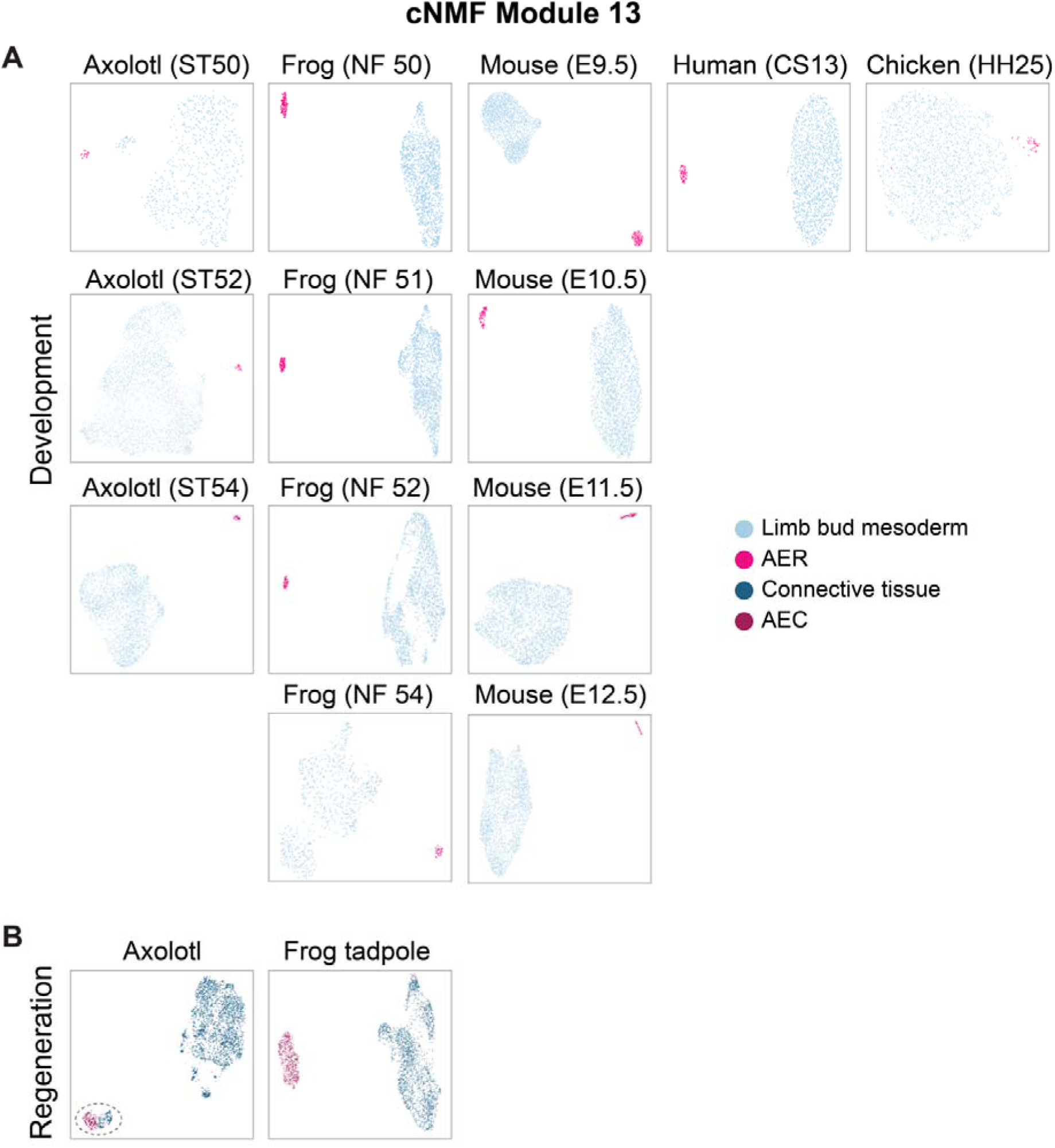
Clustering based on cNMF module 13 for individual datasets. UMAP visualization of the clustering of indicated datasets of limb development (A) and regeneration (B) based on the top 500 genes from AER-specific cNMF module 13 (Figure S14 and Supplementary Table 4). In development datasets: light blue, limb bud mesoderm cells; pink, AER cells. In regeneration datasets: dark blue, CT cells; dark pink, AEC cells. Connective tissue cells gathered with the AEC population are highlighted with a dashed line.

**Figure S20.**
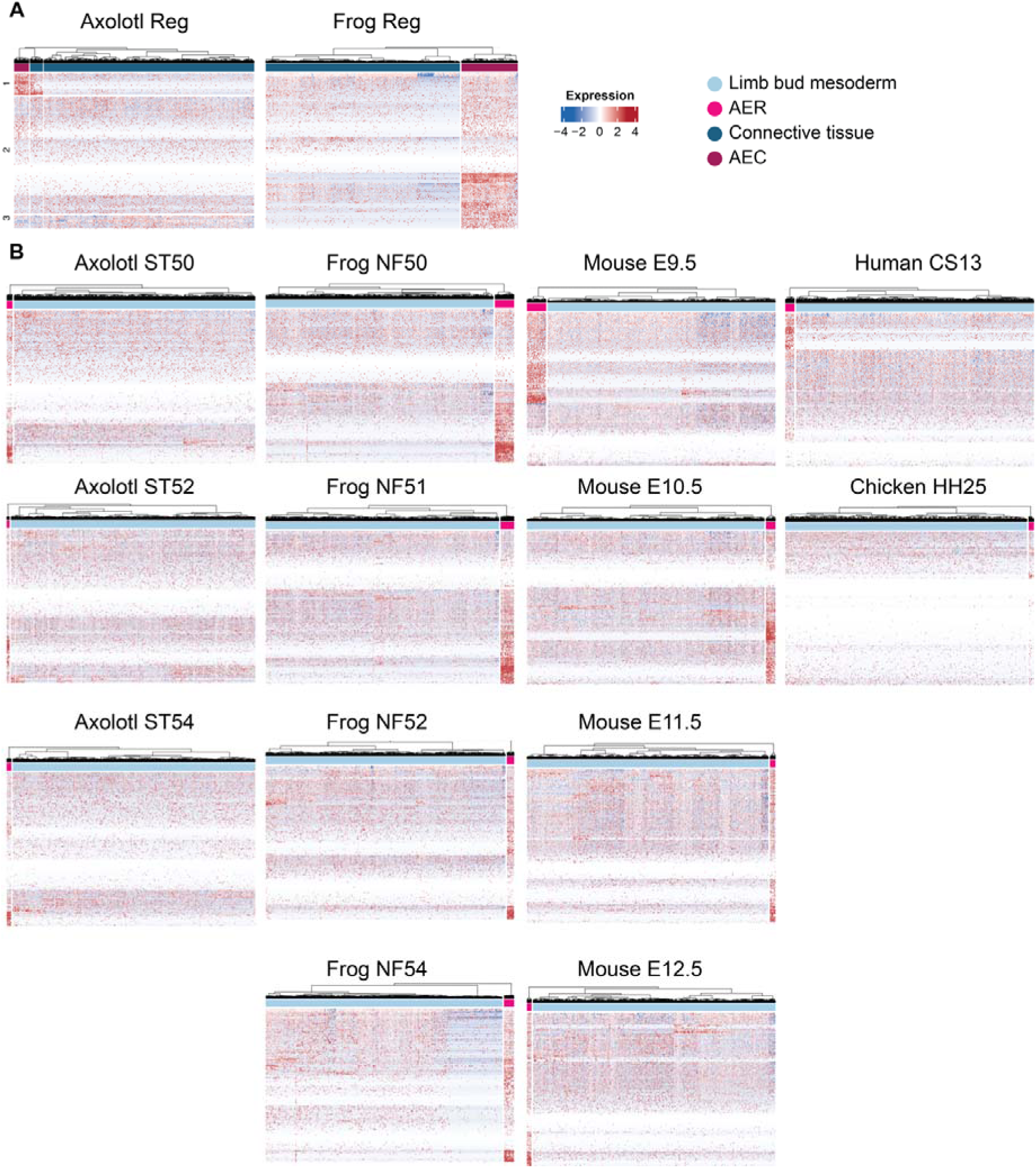
AER gene expression profile in AER, AEC, limb bud mesoderm, and connective tissue cells during limb development and regeneration. Heatmaps showing the expression profile of 500 AER DEGs in regenerating limbs (A) and developing limbs (B). For axolotl regenerating limbs (Axolotl Reg), genes were grouped into three by the K-means algorithm, and the first group is shown in Figure 3D. Please note that part of the axolotl regeneration dataset is also presented in Figure 3F. Colored squares above the heatmap represent cells from indicated populations: light blue, limb bud mesoderm cells in the development dataset; light pink, AER cells in the development dataset; dark blue, connective tissue cells in the regeneration dataset; dark pink, AEC cells in the regeneration dataset.

**Figure S21.**
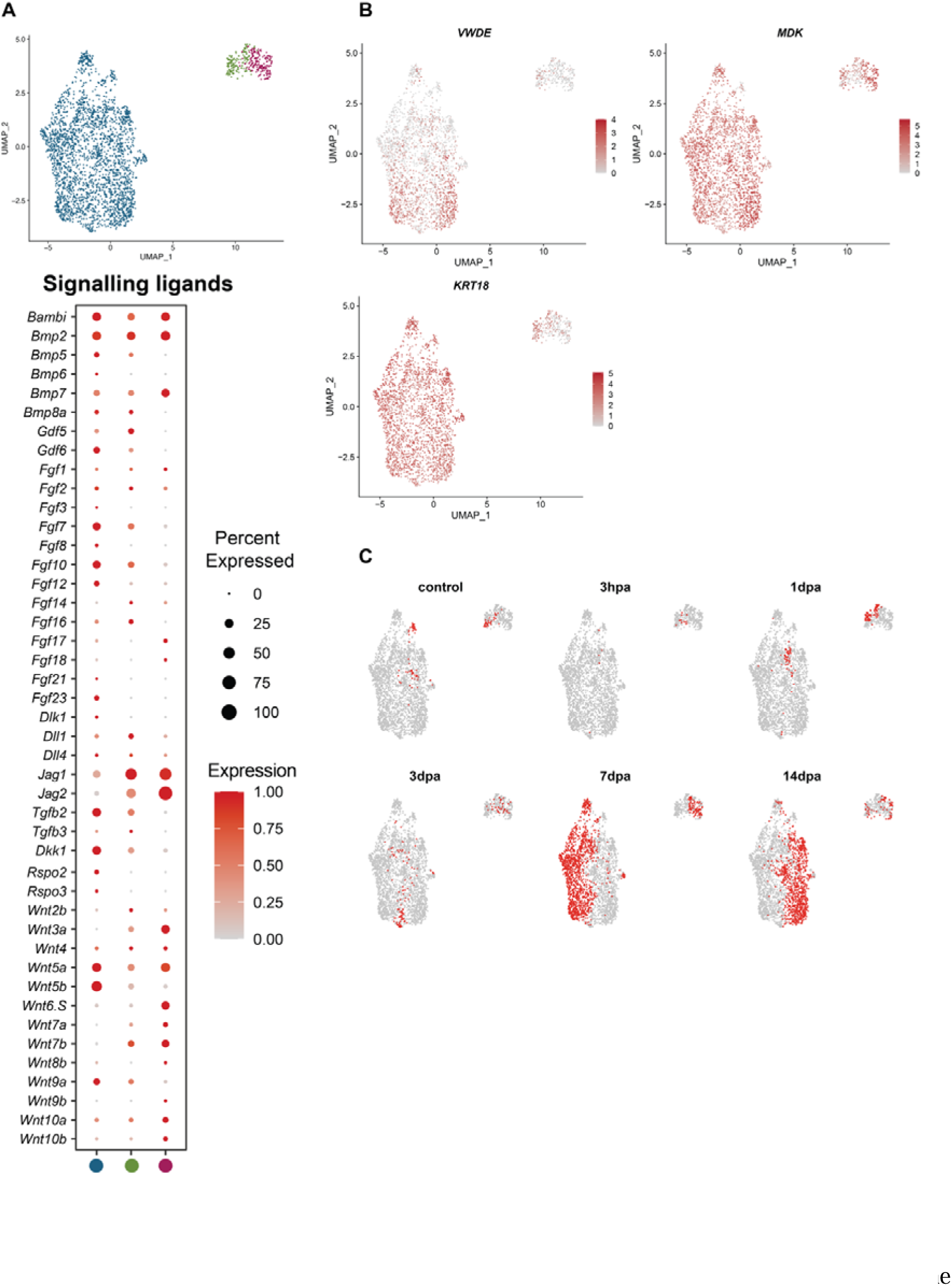
Mesodermal cells showing AER transcriptional program are present in intact limbs and express signaling ligands. A. (Top) UMAP plot from Figures 3C and S15 axolotl regeneration dataset is used to distinguish mesodermal cells showing part of the AER program and labeled by green. The rest of the connective tissue is labeled dark blue, and AEC is labeled dark pink. (Bottom) Dotplot showing signaling ligands expressions in the populations distinguished in Figure S20A. The dot color indicates the mean expression that was normalized to the max of each cell type and to the max of each gene; the dot size represents the percentage of cells with non-zero expression. B. The expression profile of previously reported regeneration-associated genes (*18*–*20*) in the UMAP plot of clustering in Figure 3C axolotl regeneration dataset. C. Sample contribution to UMAP plot of re-clustered regeneration dataset is visualized. Red dots indicate cells from the selected sample; gray dots indicate all the other cells.

